# Imaging Cellular Metabolic Rewiring with SuMMIT-SRS

**DOI:** 10.1101/2025.10.10.681769

**Authors:** Yajuan Li, Zhi Li, Yuhan Li, Kevin Rhine, Sural Ranamukhaarachchi, Shuo Qin, Hongje Jang, Jorge Villazon, Zhiliang Bai, Stephanie I. Fraley, Gene W. Yeo, Rong Fan, Lingyan Shi

**Author notes:** Correspondence: L. S.

## Abstract

Cells dynamically rewire their metabolic pathways in response to physiological and pathological cues. Such plasticity is particularly critical in neurons, stem cells, cancer cells, and immune cells, where biosynthetic demands can shift rapidly. However, current metabolic imaging techniques using isotope labeling typically track only one metabolite at a time, limiting their ability to capture the rapid dynamics of complex metabolic networks including coordinated precursor utilization, crosstalk, and turnover. Here, we present Subcellular Multiplexed Metabolic Isotope Tracing Stimulated Raman Scattering microscopy (SuMMIT-SRS), a platform that enables simultaneous visualization of multiple metabolic dynamics at subcellular resolution. By exploiting the distinct vibrational signatures of carbon–deuterium bonds derived from multiple deuterated amino acids, lipids, and monosaccharide tracers, SuMMIT-SRS maps co-regulated DNA, RNA, protein, and lipid synthesis at the same time and resolves various individual amino acid-mediated metabolic pathways within intact cells and tissues. We demonstrate SuMMIT’s broad utility across *Drosophila* fat body tissue and developing brain, tumor organoids, aged human neurons, and mouse liver, capturing cell type–specific metabolic rewiring under genetic and pathological perturbations. This approach extends SRS to multiplexed isotope tracing, offering a powerful tool to uncover dynamic and complex biosynthesis programs in development, health, and disease.

## Introduction

Metabolic rewiring is a fundamental process that enables cells and tissues to adapt to changing physiological demands, environmental stressors, and disease states ^1–5^. Rather than being a passive reflection of cellular activity, metabolic reprogramming actively shapes cellular behavior by regulating energy production, biosynthesis, and reduction-oxidation (redox) balance, as well as generating signaling metabolites that influence gene expression and epigenetic states ^5^. Emerging evidence indicates a complex interplay between metabolic rewiring and aging or disease ^6–8^. On one hand, metabolic rewiring can drive aging and disease progression by altering the cellular metabolite landscape, for example, by producing metabolites that hijack key signaling pathways ^9^, by generating metabolites and cofactors that act as agonists or antagonists of functional proteins critical for metabolic regulation, or by reshaping cellular metabolic demands to enable adaptation to diverse microenvironments ^4,10^. Conversely, aging and disease can reprogram cellular metabolism by directly altering the expression and activity of metabolic enzymes ^11^. Aged or diseased cells frequently exhibit impaired mitochondrial function ^12^ and disrupted nutrient sensing ^13^, leading to altered substrate utilization that compromises energy homeostasis and anabolic capacity ^14^. Therefore, it is crucial to reveal molecular mechanisms linking aging, disease, and metabolic rewiring, and to identify metabolic targets that can be leveraged for aging suppression and disease treatment.

To fully understand cellular metabolic rewiring, it is essential to consider not only the total abundance of metabolites but also their diversity, spatial distributions and temporal dynamics, which together dictate the underlying metabolic networks ^15–17^. Subcellular metabolic imaging encompasses the spatiotemporal tracing of key processes such as protein synthesis and degradation ^18–21^, lipogenesis and lipolysis ^22–25^, and nucleic acid metabolism ^20,24,26^. It lies at the forefront of efforts to unravel how metabolic processes are spatially organized and compartmentalized within cells ^27–29^. Each of existing metabolic imaging tools offers distinct advantages along with limitations in resolution, live-cell compatibility, and molecular specificity. Positron emission tomography (PET) ^30^ and nuclear magnetic resonance (NMR) spectroscopy ^31^ provide valuable metabolic insights but lack the spatial resolution required for organelle-level imaging. Complementary approaches such as NanoSIMS ^32^ and multi-isotope imaging mass spectrometry (MIMS) ^33,34^ enable isotopic imaging with 50–100 nm spatial resolution, but they are inherently destructive, unsuitable for live-cell studies, and often require labor-intensive sample preparation and complex image coregistration, limiting their integration with other imaging modalities. Fluorescence-based tools, including genetically encoded metabolic sensors, offer dynamic and compartment-specific readouts of metabolic activity ^35,36^. However, they rely on genetic manipulation and exhibit context-dependent performance, restricting their use in certain *in vivo* settings or primary tissue models.

Stimulated Raman Scattering (SRS) microscopy is a label-free optical technique that captures the unique vibrational information of chemical bonds in biomolecules, allowing subcellular-resolution chemical analysis within cells and tissues ^37–40^. It offers high-fidelity Raman spectra and linear concentration dependence without nonresonant background ^26^. Its spatial resolution and tissue penetration are similar to those in two-photon fluorescence microscopy, making it suitable for high-resolution imaging of animal tissues. Using stable isotope labeling, SRS microscopy has proven instrumental in uncovering metabolic dynamics in living systems. It offers a non-invasive means to visualize biosynthetic and bioenergetic activities with organelle- and compartment-level resolution, providing critical insights into metabolic processes at nanoscale ^20,41,42^. However, previous approaches have largely been restricted to a limited set of metabolites and have predominantly relied on single-probe labeling, thereby offering only a partial understanding of complex metabolic networks and global changes in metabolic pathway regulation. The ability to simultaneously label multiple metabolites, tracking metabolic activities, tracing specific pathways, and detecting metabolic rewiring has remained an unmet yet pressing need.

To address the challenge of mapping metabolic pathways at high spatial resolution, we developed Subcellular Multiplexed Metabolic Isotope Tracing (SuMMIT)-incorporated SRS microscopy. By simultaneously tracing the incorporation of multiple metabolite precursors, SuMMIT-SRS enables real-time, high-resolution visualization of macromolecular synthesis and metabolic fluxes at the single-cell level (Fig 1a and Supplementary Fig 1a). We applied this approach across a variety of biological systems, including tumor organoids, *Drosophila* body fat and developing brain, mouse liver, and aged human neurons, to investigate the spatial dynamics of multiple metabolic activities. When combined with genetic perturbations or nutrient manipulations, SuMMIT enables direct, in situ detection of context-specific metabolic rewiring, providing mechanistic insights into metabolic flexibility and homeostasis, and offering broad potential for studying aging- and disease-related metabolic dysregulation.

**Figure 1.**
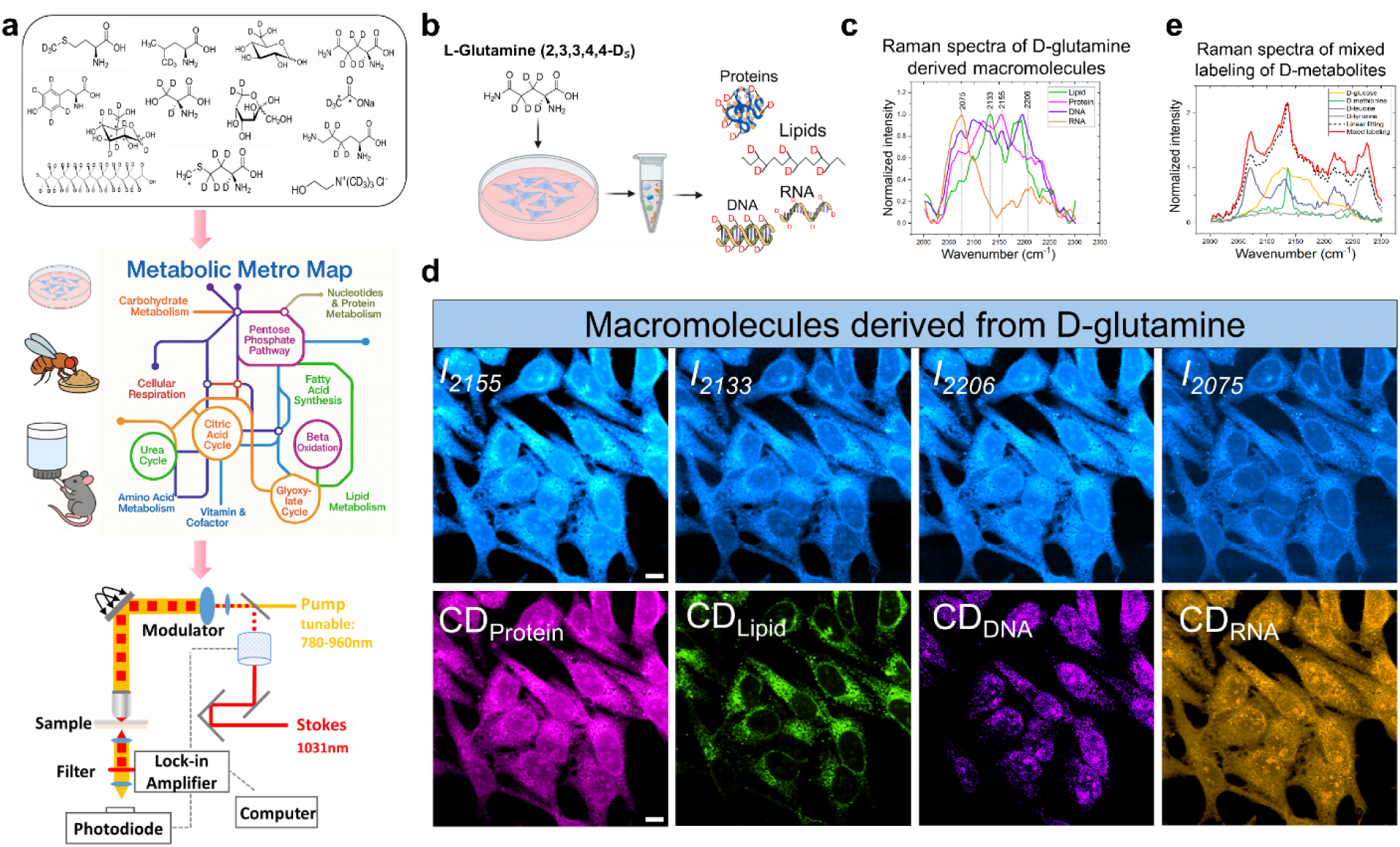
The principle of SuMMIT-SRS imaging. **(a)** Schematic showing the workflow of SuMMIT-SRS imaging. Cells, *Drosophila* flies, mice were cultured in medium or supplied with water containing deuterium labeled metabolites (D-metabolites). Following labeling, deuterium was incorporated into macromolecules including proteins, lipids, DNA, and RNA through various metabolic pathways. After that the distribution of proteins, lipids, DNA, and RNA can be captured by stimulated Raman scattering (SRS) imaging. **(b)** Cartoon showing HeLa cells cultured in medium supplemented with 4 mM D₅-glutamine for 72 h. Following labeling, proteins, lipids, DNA, and RNA were extracted from D₅-glutamine–labeled cell pellet and then subjected to Ramam spectral measurements. **(c)** Raman spectra of D₅-glutamine labeled protein, lipid, DNA, and RNA spectra measured by spontaneous Raman microscopy. **(d)** SRS images showing SuMMIT-SRS detected protein, lipid, DNA, and RNA distributions before and after the linear unmixing. Scale bar: 10 µm. **(e)** Raman spectra of HeLa cells labeled with D-glucose, D-methionine, D-tyrosine and D-leucine, respectively. They were collected using spontaneous Raman microscopy and used to generate the linear fitting spectrum. This computed spectrum agrees well with the spectrum directly measured from the cells labeled with a cocktail of D-glucose, D-methionine, D-tyrosine and D-leucine. Each spectrum was averaged of n=15 ROIs from 3 independent experiments.

### SuMMIT-SRS trace diverse origins of protein, lipid, DNA, RNA metabolism *in situ*

To map how and where diverse precursor metabolites are incorporated into macromolecular synthesis, we screened a panel of 30 deuterium-labeled metabolites (D-metabolites). These tracers allow us to image and track the incorporation of specific metabolites into proteins, lipids, nucleic acids, and other biosynthetic products across living systems (Fig 1a, Supplementary Table 1). The screening identified 18 D-metabolites exhibiting high labeling efficiency (Supplementary Fig 1a). This group of metabolic probes include various saccharide, amino acid and lipid ditopologies. To characterize the specific carbon–deuterium (C–D) Raman signatures of proteins, lipids, and nucleic acids derived from individual D-metabolites, we treated cultured cells with each D-metabolite and subsequently collected Raman spectra from purified protein, lipid, DNA, and RNA fractions. All the resulting Raman spectra revealed a distinct vibrational band around 2,000–2,300 cm^-1^ within the cell-silent region (1,800–2,600 cm^-1^), indicating the metabolic incorporation of deuterium into macromolecular structures (Supplementary Fig 1a).

We confirmed that macromolecules synthesized from D_7_-glucose and D_2_O exhibit distinguishable C–D Raman spectral signatures (Supplementary Fig 1a), as reported in other studies ^20,24^. Interestingly, proteins, lipids, and nucleic acids exhibited distinct spectral features when labeled with a range of deuterium-labeled metabolites, including D-leucine, D-methionine, D-serine, D-glutamine, D-lysine, D-choline, and D-fructose. By contrast, labeling with D-acetate, D-palmitic acid, D-oleic acid, and D-cholesterol produced signals exclusively associated with lipids (Supplementary Fig 1a). Importantly, each metabolite does not necessarily contribute to all the macromolecular synthesis pathways under investigation but indeed shows specificity in its incorporation to distinct macromolecular products. The results underscore the molecular specificity of C–D vibrational profiles and demonstrate how different metabolic precursors selectively contribute to distinct classes of macromolecules. In essence, a single D-labeled precursor can facilitate multiplexed imaging of its downstream metabolic products, capture the trajectory of its incorporation and elucidate the metabolic pathways it engages in.

We utilized D_5_-glutamine as an example to demonstrate the macromolecule-specific nature of C–D Raman signals arising from individual D-metabolites (Fig 1b-d and Supplementary Fig 1b). HeLa cells were cultured in medium supplemented with 4 mM D_5_-glutamine for 72 h, which fully replaced the standard exogenous glutamine in the culture medium (Fig 1b). A broad C–D vibrational band spanning 2,050–2,250 cm^-1^ within the Raman-silent region was detected from the resulting cell pellet. This signal was absent in unlabeled control cells, indicating the success of deuterium incorporation into newly synthesized macromolecules (Supplementary Fig 1b). During the whole labeling process, no observable toxicity or cell death was detected. Notably, each macromolecular fraction derived from D₅-glutamine exhibited a distinct C–D spectral profile, with unique peak positions and shapes. Proteins and DNA displayed complex spectral features with multiple peaks centered around 2,075, 2,110, 2,155, and 2,187 cm^-1^ (Fig 1c and Supplementary Fig 1b), while RNA and lipids showed simpler spectra with only one or two dominant peaks, respectively (Fig 1c and Supplementary Fig 1b). The divergence in proteins, lipids, DNA, and RNA spectral profiles reflects the atom-resolved tracing of deuterium through central metabolic pathways. More specifically, the deuterium atoms of D₅-glutamine are incorporated into proteins (via glutamine transamination to glutamate and subsequent entry into the amino acid pool), nucleotides (via anaplerosis and TCA cycle), and lipids (through TCA cycle carbon flux contributing to acetyl-CoA production) ^43^.

Linear spectral fitting of the Raman signals from isolated cellular proteins, lipids, DNA, and RNA showed strong agreement with the overall spectrum of the D₅-glutamine–labeled cell pellet (Supplementary Fig 1b), indicating that these four macromolecule classes account for the majority of the observed C–D stretching vibrations. Due to the spectral overlap of vibrational bands from proteins, lipids and nucleic acids, SRS imaging acquired at any single frequency of the C–D region actually involves a mixed signal of D₅-glutamine–derived C–D bonds across various biomolecular pools. By applying a linear decomposition algorithm, such an overlapping SRS signal can be decomposed into macromolecule-specific components. The underlying principle is that for *n* chemical species with distinguishable Raman spectra, linear unmixing can be performed using signal intensities at *n* distinct frequencies (or channels). Based on this, we acquired SRS images at selected Raman shifts (2,076, 2,133, 2,155, and 2,206 cm⁻¹) by tuning the laser wavelength accordingly (Fig 1d). Normalized reference spectra from standard compounds representing lipids, proteins, DNA, and RNA were used to calculate the unmixing coefficients (Supplementary Fig 1c), which were then applied to reconstruct the unmixed images. Leveraging the strong correspondence between SRS and Raman spectra, along with the linear relationship between SRS intensity and molecular concentration ^39,44^, we successfully mapped the spatial distribution of each D₅-glutamine–derived macromolecule at the cellular and subcellular level via SRS imaging (Fig 1d).

The strategy is broadly applicable to a range of deuterium-labeled metabolic probes. As shown in (Supplementary Fig 1a), in multiple cases, different macromolecules derived from the same D-metabolite exhibit distinct spectral signatures for proteins, lipids and nucleic acids, reflecting the unique chemical environments of their C–D bonds. By harnessing the spectral specificity of C–D vibrations, this approach enables simultaneous in situ visualization of D-labeled lipids, proteins, DNA, and RNA, providing insights into how each D-metabolic probe is selectively routed into distinct biosynthetic pathways.

Inspired by the observation that each macromolecule displays distinct spectral signatures depending on its D-labeled precursors, we hypothesized that it may be feasible to resolve the synthetic activity and spatial distribution of multiple D-metabolite–derived macromolecules simultaneously within the same biological sample. Therefore, we labeled cells with a combination of D-glucose, D-methionine, D-leucine, and D-tyrosine. Linear fitting of the four Raman spectra from singly labeled cells showed strong agreement with the spectrum obtained from the pellet of cells treated with combination labeling (Fig1e), indicating that the specific C–D Raman fingerprints of each precursor are well preserved under mixed labeling conditions. These results validate the feasibility of spectral unmixing to resolve the individual contributions of various metabolic precursors in complex biological contexts. When combined with SRS microscopy, this approach enables accurate and multiplexed mapping of macromolecular synthesis from diverse precursors –– a strategy we term Subcellular Multiplexed Metabolic Isotope Tracing (SuMMIT) SRS imaging.

### SUMMIT-SRS reveals the metabolic heterogeneity in breast cancer organoids

We next applied SuMMIT-SRS imaging to 3D cultured breast cancer (BRCA) organoids that spontaneously gives rise to two distinct and stable morphologies, collectively invasive networks and compact, non-invasive spheroids— thereby recapitulating the architectural and functional heterogeneity observed in vivo (Fig 2a) ^45,46^. To explore potential differences in metabolic rewiring between these phenotypes and gain mechanistic insights into cancer metabolism, we performed combinatorial labeling with organoids using D-glucose, D-glutamine, D-methionine, D-leucine, and D-tyrosine. This combinatorial labeling covers major classes of macromolecules (lipids, proteins, nucleic acids) while also offering spectrally distinguishable C–D signatures. Following labeling, we extracted pure proteins, lipids, and DNA from labeled BRCA cells and verified deuterium incorporation into each macromolecule under this labeling condition (Supplementary Fig 2a and 2b).

**Figure 2.**
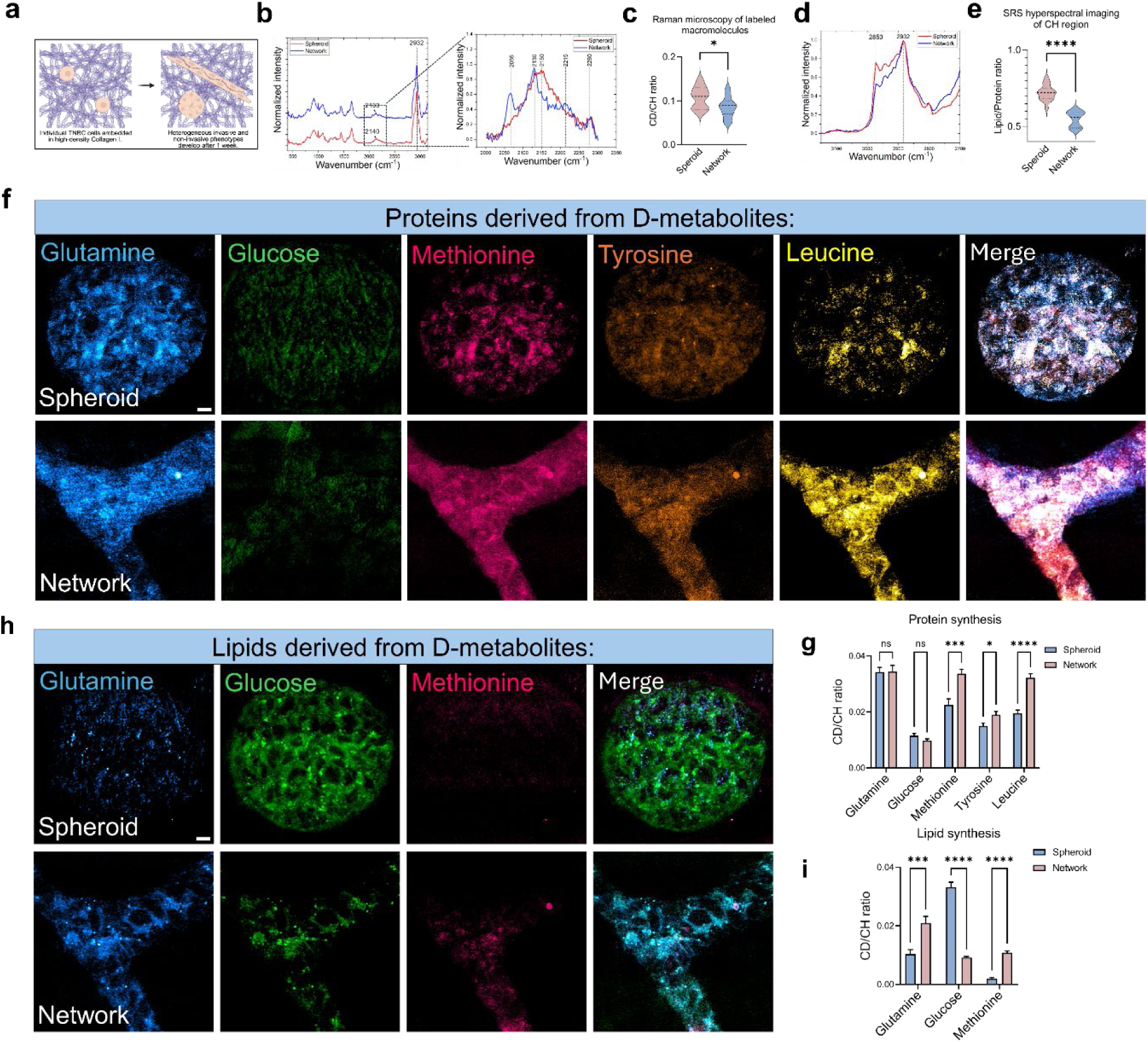
SuMMIT-SRS imaging reveals the metabolic heterogeneity in the breast cancer model. **(a)** Schematic of the invasive (networks) and noninvasive (spheroids) tumor organoid model. **(b)** Raman spectra of tumor networks and spheroids after 72 h of simultaneous co-labeling with D-glucose, D-glutamine, D-methionine, D-leucine, and D-tyrosine. Inset shows a zoomed-in view of the C–D region. **(c)** Quantification of macromolecule synthesis using the ratio of CD to CH area under curve (AUC) based on the Raman spectra in (B). n=15 regions of interest (ROIs) per group. **(d)** Raman spectra of the CH region of cytoplasm in the cells from spheroids and networks, collected by SRS-hyperspectral imaging (HSI) showing total lipid peak at 2850 cm^-1^ and protein peak at 2932 cm^-1^. Each spectrum was averaged from n=15 regions of interest (ROIs) per group. **(e)** Quantification of the lipid to protein ratio (2850 cm^-1^/3012 cm^-1^) based on the Raman spectra in (d). n=15 ROIs per group. **(f)** SRS images showing SuMMIT-SRS imaging detected single and merged channels of glutamine, glucose, methionine, tyrosine, leucine, derived protein signal in spheroids and networks. Scale bar: 10 µm. **(g)** Quantification of protein synthesis from each deuterium-labeled precursor using the CD/CH ratio based on the unmixed SRS signal in (f). n=15 ROIs per group. **(h)** SRS images showing SuMMIT-SRS-detected single and merged channels of glutamine-, glucose-, and methionine-derived lipid signals in spheroids and networks. Scale bar: 10 µm. **(i)** Quantification of the lipid synthesis from each deuterium-labeled precursor using the CD/CH ratio based on the unmixed SRS signal in (h). n=15 ROIs per group. Values in (g, i) are mean ± SEM. ns, not significant; *, p < 0.05; **, p < 0.01; ***, p < 0.001; ****, p < 0.0001 by Student’s *t*-test.

We then acquired *in situ* Raman spectra from both tumor networks and spheroids and observed a marked difference in the shape of the C–D band within the cell-silent region (Fig 2b), indicating a divergent substrate preference underlying their anabolic pathways. Based on Raman spectra, the overall anabolic activity quantified by calculating the ratio of the C–D area under the curve (AUC) to the C–H AUC was significantly reduced in the networks (Fig 2c). This result was confirmed by the greater intensity ratio between 2,130 cm^-1^ (an overall C–D peak representing the total metabolic activity) and 2,932 cm^-1^ (the highest intensity peak at CH region representing protein-related CH_3_ stretching) in spheroids (Supplementary Fig 2c). To further determine which macromolecules contribute the most to the C–D band shape differences, we performed multichannel SRS imaging and unmixing of tumor networks and spheroids at 2,066, 2,130, and 2,150 cm^-1^ guided by reference spectra of extracted protein, lipid, and DNA (Supplementary Fig 2b). The signals of newly synthesized proteins (CD_P_), lipids (CD_L_), and DNA (CD_D_) were then unmixed and quantified by C–D/C–H (CD/CH) (Supplementary Fig 2d). The quantitative analysis revealed that, compared to the non-invasive spheroids, the invasive networks exhibited a significant increase in overall protein synthesis, a decrease in overall lipid synthesis, but no significant change in DNA synthesis (Supplementary Fig 2e-g). Intriguingly, we noticed that the CD_D_ signal was more diffusely distributed throughout the cytoplasm in the networks, indicating impaired nuclear retention or mislocalization of newly synthesized DNA. Consistently, SRS hyperspectral imaging (HSI) of the C–H region demonstrated higher lipid content in spheroids, as indicated by an increased lipid-to-protein ratio (2,850/2,932 cm⁻¹) (Fig 2d and 2e). Conversely, the total protein level was lower in spheroids, as evidenced by reduced 2,932 cm⁻¹ peak intensity after lipid removal (Supplementary Fig 2h). The observed changes in protein and lipid metabolism were consistent with our prior RNA-seq analysis, which revealed upregulation of genes involved in protein biogenesis and metabolism, accompanied by downregulation of genes associated with lipid metabolism in the network phenotype ^46,47^.

We next compared the Raman spectra of purified proteins and lipids from each phenotype and observed markedly distinct features (Supplementary Fig 2i and 2j). By applying SuMMIT-SRS imaging to spatially map protein signals, we observed significant enrichment of methionine-, tyrosine-, and leucine-derived proteins in the network phenotype (Fig 2f and 2g), while protein synthesis from glutamine and glucose showed no significant difference relative to spheroids (Fig 2f and 2g). Notably, the D-labeled protein signal derived from glucose was lower than that from amino acid tracers in both phenotypes (Fig 2f and 2g), suggesting that deuterium from glucose was preferentially directed toward lipid synthesis or dissipated during metabolic processing. Consistently, RNA-seq analysis revealed upregulation of methionine and leucine metabolic pathways, alongside downregulation of the glycolytic pathway (Supplementary Fig 2l).

SuMMIT-SRS imaging of lipid synthesis demonstrated that, although total lipid production was reduced in the tumor networks, there was a pronounced rewiring of lipid biosynthetic pathways (Fig 2h, 2i and Supplementary Fig 2j). In spheroids, lipids were predominantly synthesized from glucose, with a minor contribution from glutamine, whereas methionine-derived labeling was barely detectable (Fig 2h and 2i). In contrast, lipid synthesis in the network phenotype was predominantly fueled by glutamine, whereas the contribution from glucose was reduced (Fig 2h and 2i). This observation is consistent with the reduced NADH/FAD redox ratio in networks (Supplementary Fig 2d and 2k), indicating decreased glycolytic NADH generation and a diminished supply of glycolysis-derived precursors for lipid synthesis. Notably, methionine made a greater contribution to lipid synthesis in the network phenotype (Fig 2h and 2i), indicating its diversion from protein synthesis and suggesting that transmethylation reactions or one-carbon metabolism actively support lipid biosynthesis ^26,48^. Consistently, RNA-seq analysis revealed upregulation of methionine and leucine metabolic pathways, alongside downregulation of the glycolytic pathway (Supplementary Fig 2l). Together, these results indicate that the invasive network is characterized by elevated protein synthesis, reduced lipid synthesis, and differential utilization of D-metabolite precursors for macromolecule production, reflecting distinct metabolic reprogramming in tumor phenotypes.

### SuMMIT-SRS unravels spatial metabolic reprogramming under genetic manipulation

To assess the applicability of SuMMIT for *in vivo* mapping of metabolic dynamics, we employed *Drosophila melanogaster* as a model system. *Drosophila* first instar larvae were maintained on a holidic diet ^49^ supplemented with deuterium-labeled metabolites at the physiological level for three (mid-third instar) or four (late-third instar) days (Fig 3a). Raman spectra were subsequently acquired from purified proteins, lipids, and nucleic acids from fly tissues. The resulting profiles confirmed that the Raman signatures of individual metabolite-derived macromolecules were preserved in *Drosophila*, closely resembling those observed in cultured cells (Supplementary Fig 1a). These findings indicate robust metabolite incorporation and processing in whole animal, highlighting a high degree of conservation in core metabolic pathways between *Drosophila* and mammalian systems. Using C–D spectra from fly tissues as references for linear unmixing, we successfully resolved and visualized protein and lipid signals derived from each D-metabolite in the *Drosophila* larval fat body and brain tissues (Supplementary Fig 3a).

**Figure 3.**
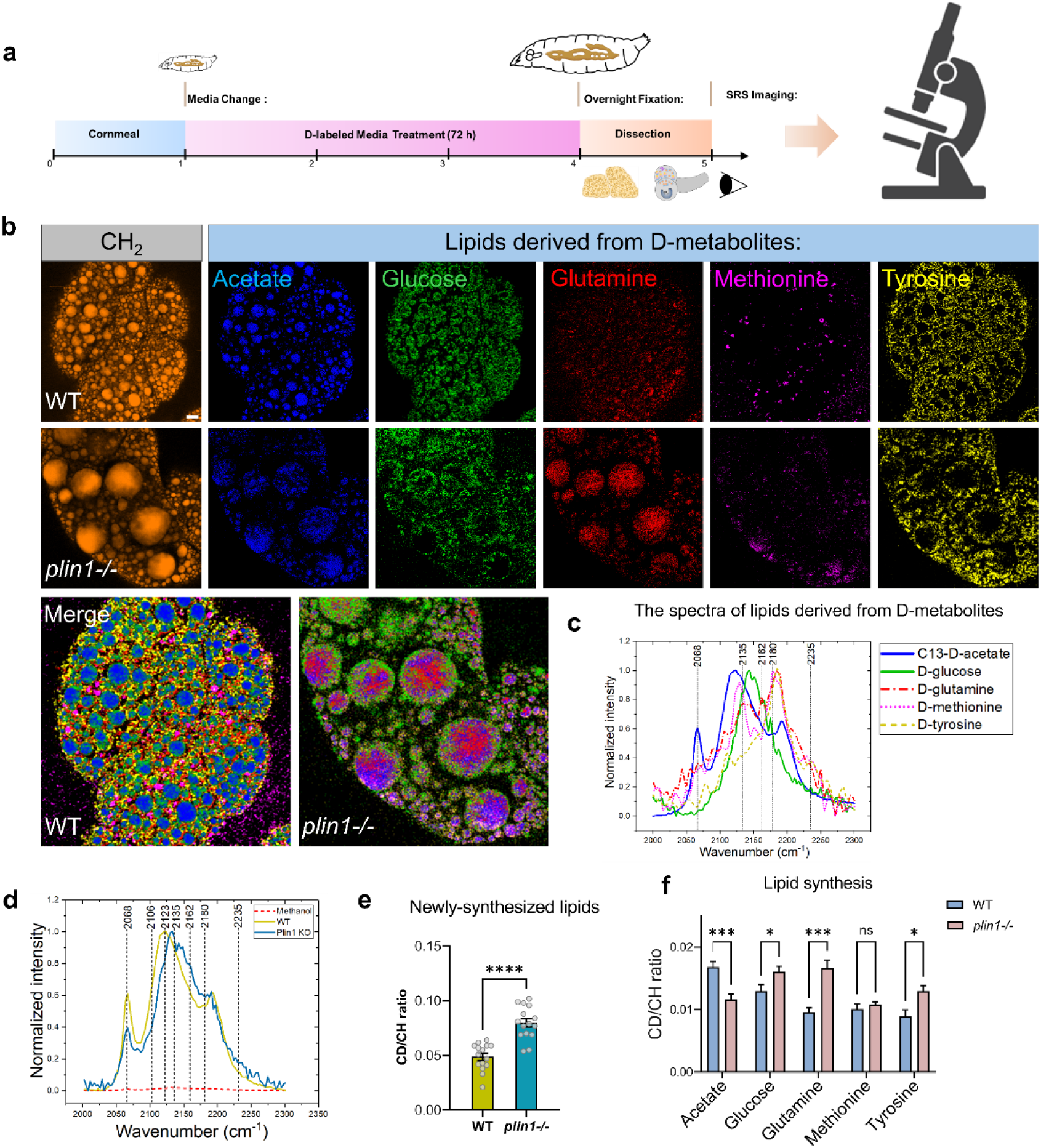
SuMMIT-SRS unravels spatial metabolic reprogramming under genetic manipulation. **(a)** Schematic depicting the labeling paradigm of *Drosophila* larvae, 1^st^ instar larvae (24 h after egg laying) were fed with labeling food for 72 h or 96 h, reaching mid-3^rd^ instar or late-3^rd^ instar stage. **(b)** Images showing SuMMIT-SRS-detected single and merged channels of acetate-, glucose-, glutamine-, methionine-, and tyrosine-derived lipid signals in the WT and Plin1 mutant fat body of *Drosophila* late third instar larvae. Scale bar: 10 µm. **(c)** Raman spectra of lipids derived from individual labeling of acetate, glutamine, glutamine, glutamine, glucose, methionine and tyrosine in the fat body of late third instar *Drosophila* larvae, measured by spontaneous Raman microscopy. Each spectrum was averaged from n=15 larvae per group. **(d)** Zoomed-in CD Raman spectra of lipids described from mix labeling with acetate, glutamine, glutamine, glutamine, glucose, methionine and tyrosine in the fat body of late third instar *Drosophila* larvae, measured by spontaneous Raman microscopy with or without methanol wash. Each spectrum was averaged from n=15 larvae per group. **(e)** Quantification of lipid synthesis by the ratio of CD (AUC) to CH (AUC) based on the Raman spectra. n=15 regions of interest (ROIs) per group. ****, p < 0.0001 by Student’s *t*-test. **(f)** Quantification of lipid synthesis from each deuterium labeled precursor by the CD/CH ratio based on the unmixed SRS signal in (b). n=15 ROIs per group. Values in (e, f) are mean ± SEM. ns, not significant; *, p < 0.05; ***, p < 0.001; ****, p < 0.0001 by Student’s *t*-test.

The fat body in *Drosophila* is a multifunctional tissue that integrates roles analogous to the mammalian liver, adipose tissue, and innate immune system ^50^. Within this tissue, lipid droplets (LDs) are abundant organelles that govern fat storage and systemic metabolic balance. The evolutionarily conserved PERILIPIN family protein Plin1 localizes to LD surfaces and serves as a key regulator of lipid homeostasis. Loss of *plin1* transforms LD morphology from multilocular to unilocular fat cells in larvae, while the total lipid content of *plin1* mutants remains comparable to that of controls ^51^.

To investigate the metabolic reprogramming associated with Plin1 deficiency, we applied SuMMIT-SRS imaging to profile macromolecular synthesis and pathway-specific dynamics in the *Drosophila* fat body. Flies were labeled with a combination of D-acetate, D-glucose, D-glutamine, D-methionine, and D-tyrosine, a panel that collectively spans complementary entry points into central carbon metabolism and macromolecular synthesis (Fig 3b and 3c). As shown in (Fig 3b), a single large LD (∼18 µm in average) remained prominently localized within the fat body cells of *plin1* knockout larvae following labeling. This observation indicates that the administration of D-metabolites did not disrupt the characteristic LD phenotype. Raman spectrometry detected distinct Raman shapes in wild-type (WT) and *plin1* mutant flies (Fig 3d and Supplementary Fig 4a) and the newly synthesized lipids are increased in *plin1* mutant as quantified by CD/CH (Fig 3e). All D-metabolite–derived signals on LDs were abolished by methanol washing, confirming their association with lipid species (Fig 3d).

Following multichannel imaging (Fig 3b and Supplementary Fig 4b) and spectral unmixing (Fig 3c and Supplementary Fig 4c), signals from *de novo* lipid synthesis derived from each D-labeled tracer were comparable in intensity and could be successfully separated. Each tracer revealed distinct spatial distribution patterns, indicative of substrate-specific lipid synthesis and compartmental routing. In both WT and *plin1* mutant fat bodies, acetate-derived lipids were consistently concentrated in the core of LDs, whereas glucose-derived lipids exhibited an uneven patch-like distribution, suggesting potential organelle-organelle contact sites, likely corresponding to regions of close association between LDs and the endoplasmic reticulum (ER)—the primary site of lipogenesis. D-glutamine– derived signals were predominantly localized to small LDs in WT fat bodies, while both large and small LDs showed incorporation of D-glutamine–derived signals in *plin1* mutants (Fig 3b), suggesting active glutamine-driven lipogenesis across LD populations. For D-tyrosine, signals in WT fat bodies were primarily localized to the periphery (Fig 3b), potentially associated with membrane-rich structures. In contrast, in the *plin1* mutant, a small but distinct portion of the D-tyrosine signal was redirected to LDs, suggesting a shift in tyrosine metabolism toward lipid storage rather than structural or signaling functions under metabolically reprogrammed conditions.

Unexpectedly, D-methionine–derived signals showed strong co-localization with lysosomes in WT flies (Fig 3b and Supplementary Fig 4d), as confirmed by Lysotracker staining (Supplementary Fig 4d). This lysosomal enrichment was abolished in *Atg5* knockdown animals (Supplementary Fig 4d), implicating autophagy as a critical pathway mediating the trafficking or turnover of methionine-derived lipids. Compared with WT flies, the lysosomal localization was absent in *plin1* mutants. Instead, methionine-derived signals are partially localized to the LD core, suggesting a shift in lipid routing or turnover (Fig 3b). These findings reveal a previously unappreciated developmental and spatial plasticity in methionine metabolism, linking it to organelle-specific lipid handling via autophagy.

Notably, we found D-methionine labeling yielded five distinct Raman spectral shapes throughout *Drosophila* fat body development (Supplementary Fig 4e). A unique spectral feature was observed in early-stage larvae, characterized by a single C–D peak at 2140 cm⁻¹ (Supplementary Fig 4e). SRS imaging at this wavenumber revealed strong D-methionine–derived signals in LDs (Supplementary Fig 3a). In contrast, other developmental stages typically exhibited a broader spectrum of D-methionine-labeled signals with two distinct peaks (Supplementary Fig 4e). Intriguingly, the Raman spectra from mature adults (72 h after eclosion) closely resembled those of late third-instar larvae (before metamorphosis) (Supplementary Fig 4e), indicating a conserved metabolic fate of methionine at these stages. However, during the pupal stage, a dynamic shift in the intensity of the 2185 cm⁻¹ peak was observed (Supplementary Fig 4e), which is predominantly attributed to D-labeled lipids in LDs (Supplementary Fig 4f and 4g). This lipid-specific signal was confirmed by its loss following methanol wash and its retention after proteinase K treatment (Supplementary Fig 4g).

Furthermore, subcellular localization of the D-methionine signal also changed during development. In late-stage larvae, D-methionine–derived signal localized primarily to lysosomes (Fig 3b and Supplementary Fig 4d), while in mature adults, the signal was predominantly associated with LDs (Supplementary Fig 4g)—despite similar Raman spectral shapes (Supplementary Fig 4e). Collectively, these findings reveal the distinct metabolic routing of precursor substrates and suggest that *plin1* deficiency leads to a remarkable metabolic reprogramming at late third instar larvae, including altered lipid synthesis pathways, precursor utilization, and subcellular lipid trafficking.

In addition to providing spatial information, SuMMIT-SRS enables quantitative analysis by calculating the intensity ratio of C–D to C–H lipid channels, which serves as a proxy for the renewal and turnover of lipids derived from specific precursors. In *plin1* mutants, we observed a remarkable increase in glucose- and glutamine-derived lipids alongside a decrease in acetate-derived lipids, suggesting a compensatory mechanism to maintain overall lipid homeostasis (Fig 3b and 3f). Furthermore, relative ratios between different macromolecular channels were readily visualized in the overlaid absolute-intensity images (Fig 3b), demonstrating the quantitative reliability of SuMMIT-SRS for assessing metabolic heterogeneity at the subcellular level.

To enhance data interpretation and throughput of the SuMMIT-SRS system, we developed Computational Mapping of Predicted Active Subcellular Substrates (COMPASS), a deep learning pipeline for automated detection and quantification of metabolite distributions within complex tissue environments (Fig 4a). COMPASS utilizes a one-dimensional residual neural network (1D-ResNet) architecture to predict the spatial distribution of various isotope-labeled metabolites from SRS hyperspectral Raman images (HSI). The framework first extracts feature representations from reference C–D spectral data through feature extraction and classification learning (Supplementary Fig 4h), then applies these learned features (Supplementary Fig 4i) to C–D hyperspectral images for pixel-wise prediction.

**Figure 4.**
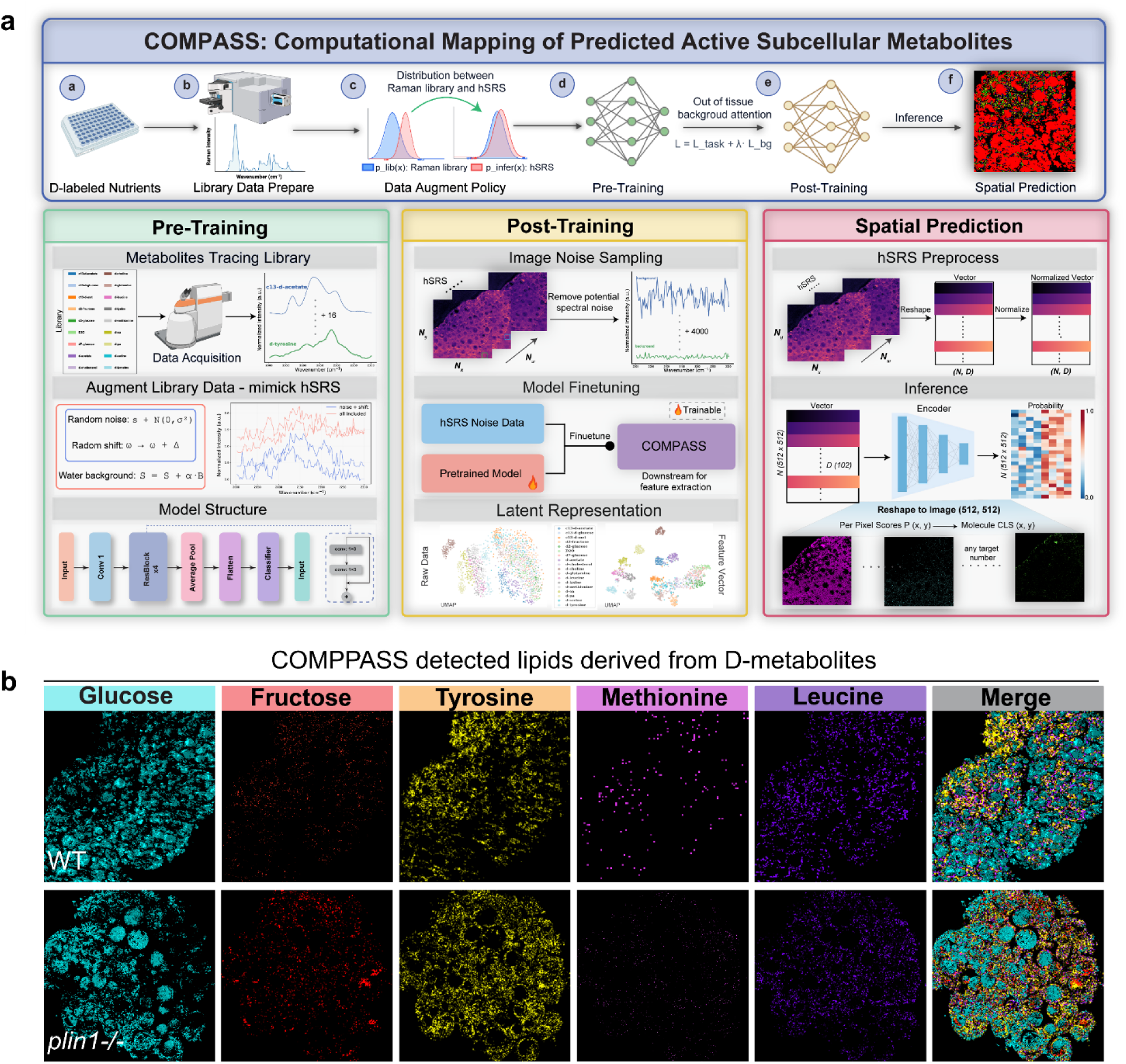
COMPASS for automated detection and quantification of metabolic distribution. **(a)** COMPASS computational framework for metabolite prediction in hyperspectral SRS images. The workflow consists of (a-e) D-labeled nutrient acquisition, Raman spectral collection, neural network training, fine-tuning, and hyperspectral SRS (hSRS) inference. The three-stage implementation includes: pretraining on reference spectra with data augmentation strategies to mimic real-world conditions; post-training with real hSRS noise integration for model fine-tuning; and inference on hyperspectral images for spatial metabolite mapping. Model interpretability is enhanced through feature visualization and Grad-CAM, enabling identification of spectral regions critical for metabolite prediction.**(b)** COMPASS-detected single and merged channels of glucose-, fructose-, glutamine-, methionine-, tyrosine-, and leucine-derived lipid signals in the fat body of WT and Plin1 mutant late third instar *Drosophila* larvae. Scale bar: 10 µm.

Our model effectively mapped the spatial distribution of diverse macromolecules derived from mixed deuterium labeling of D-methionine, D-tyrosine, D-leucine, and even distinguished subtle differences between D-glucose- and D-fructose-derived metabolites within *Drosophila* fat bodies (Fig 4b). Importantly, the findings from both linear spectral unmixing and AI detection were further validated by direct C–D/C–H quantification in single-metabolite labeling experiments (Supplementary Fig 4j). All these data together demonstrate that the integration of SuMMIT with COMPASS offers a powerful platform for quantitative imaging of multiple metabolic pathway and enables efficient detection of metabolic rewiring across different physiological and developmental contexts.

### SuMMIT-SRS unveils carbon source reprogramming for hepatic *de novo* lipogenesis under intermittent fasting

Functionally analogous to the *Drosophila* fat body, the mammalian liver exhibits dynamic metabolic remodeling in response to various stresses and pathophysiological states, such as intermittent fasting. However, how to spatiotemporally assess hepatic metabolic flexibility at subcellular resolution remains technically challenging.

We applied SuMMIT-SRS to trace metabolite precursor utilization in mouse liver. In mice, deuterium-labeled metabolite tracing via drinking water offers a convenient and non-invasive strategy for assessing metabolic flux *in vivo* (Fig 5a) ^52^. In our labeling experiment, the control group received continuous administration of D-glutamine (2.5 mg/mL), D-acetate (10 mg/mL), and D-methionine (2.5 mg/mL) in drinking water for 10 consecutive days (Supplementary Fig 5a). This combination was selected based on prior mass spectrometry analyses, which showed that amino acids contribute to *de novo* lipogenesis (DNL) with approximately 14- and 8-fold greater efficiency than glucose ^52^. In parallel, acetate has been identified as a particularly prominent lipogenic substrate in the liver ^52^ .

**Figure 5.**
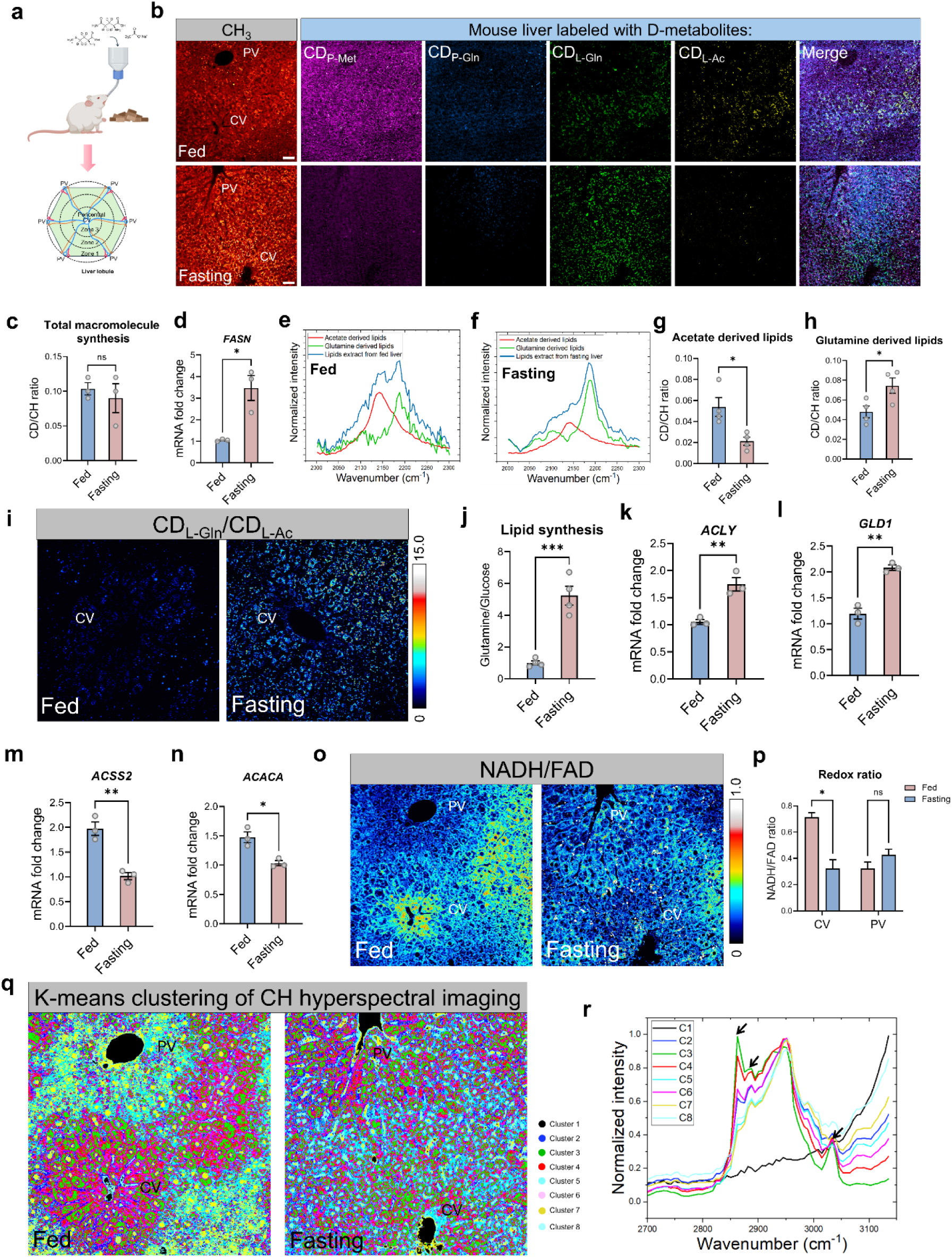
SuMMIT-SRS imaging dissects carbon source for hepatic *de novo* lipogenesis under intermittent fasting. **(a)** Cartoon showing the mice fed with D-labels in drinking water with the liver sections examined by SuMMIT-SRS imaging. **(b)** Images showing SuMMIT-SRS-detected single and merged channels of acetate-, glutamine-, and methionine-derived protein and lipid signals in the fed and fasting mouse liver. Scale bar: 20 µm. **(c)** Quantification of macromolecule synthesis by the ratio of CD (AUC) to CH (AUC) based on the Raman spectra. n=3 mice per group. **(d)** qRT-PCR showing an increased expression of *FASN* gene in fasting liver compared to the control group. n=3 mice per group. **(e-f)** Raman spectra of the CD region of acetate-, glutamine-, and mix labeling-derived lipids from fed(e) and fasting(f) liver, respectively. Each spectrum was averaged from n=30 spectra collected from 4 mice per group. **(g-h)** Quantification of lipid synthesis from D-acetate (g) and D-glutamine (h) by the CD/CH ratio based on the unmixed SRS signals. n=4 mice per group. **(i)** SRS ratiometric images generated by the intensity ratio of D-glutamine derived lipids to D-acetate derived lipids based on the unmixed SRS signal. n=4 mice per group. **(j)** Quantification of the intensity ratio of D-glutamine derived lipids to D-acetate derived lipids based on (i). n=4 mice per group. **(k-n)** qRT-PCR showing an increased expression of genes in glutamine metabolic pathway (*ACLY* (k) and *GLD1* (l)) but a reduction of gene expression in acetate metabolic pathway (*ACSS2* (m) and *ACACA* (n)) in fasting liver compared to the control group. n=3 mice per group. **(o)** Redox ratio images (NADH/FAD) generated by two photon excited fluorescence (TPEF) imaging detected NADH and FAD signal in the fed and fasting liver. **(p)** Quantification of redox ratio within PV and CV regions in the fed and fasting liver. n=30 ROIs from 3 mice for each group. **(q-r)** Stimulated Raman Scattering Hyperspectral Imaging (SRS-HSI) combined with unsupervised K-means clustering applied to the C–H stretching spectral region to analyze liver tissue under fed and fasting conditions. Each cluster, identified by a distinct color, corresponds to a unique chemical profile derived from the spectral data. The spatial localization of each color in the image reveals the distribution of these chemical features across the tissue (q). The spectral plot displays the average Raman spectra for each cluster (r), with the curve color matching the corresponding spatial region, thereby enabling direct correlation between chemical composition and tissue morphology. C, cluster. The arrows indicate lipid peaks at 2850, 2880, and 3015 cm⁻¹. Each clustering image represents 9 ROIs from 3 mice for each group. Values in (c, d, g, h, j-n, p) are mean ± SEM. ns, not significant; *, p < 0.05; **, p < 0.01; ***, p < 0.001 by Student’s *t*-test.

Following metabolic labeling, liver tissue sections collected after sacrifice were analyzed using spontaneous Raman microscopy. A strong C–D signal was detected, confirming effective incorporation of the labeled precursors under the administered labeling conditions (Supplementary Fig 5b). To determine the macromolecular sources of the signal, we biochemically isolated pure fractions of proteins, lipids, and DNA from the tissue. The C–D signal was detectable in each of these macromolecular fractions (Supplementary Fig 5c). By using SuMMIT-SRS, we successfully mapped the distributions of newly synthesized proteins, lipids, and DNA within the liver tissue (Supplementary Fig 5d). Notably, the spatial distributions of C–D labeled proteins, lipids, and DNA aligned well with the CH signals of proteins, lipids, and DNA collected at 2,967, 2,926, and 2,850 cm^-1^ as in previous report ^26^, supporting the fidelity of our metabolic mapping approach.

To examine whether amino acids and acetate metabolic fluxes in hepatocyte at different regions were altered during intermittent fasting, the experimental group of mice was subjected to 2:1 (2 days of feeding, 1 day of fasting) pattern intermittent fasting, during which they were provided with access to drinking water containing the same concentration of D-labeled metabolites as the control group. During feeding phases, the mice were given normal chow diet. The entire treatment lasted for 10 days (Supplementary Fig 5a). By SuMMIT-SRS, we observed a greater LD accumulation in the fasting liver (Fig 5b), although the total macromolecule synthesis had no significant difference between control and fasting group (Fig 5c). Consistent with LD accumulation, quantitative reverse transcription PCR (RT-qPCR) showed a significant increase of mRNA levels of *FASN*, which is involved in lipid synthesis in the fasting liver (Fig 5d). Furthermore, SuMMIT imaging-based metabolic pathway analysis revealed a reduction in protein synthesis from methionine and glutamine (Fig 5b, Supplementary Fig 5e and 5f) while a rewiring of lipid synthesis from divergent precursors in the intermittent fasting liver compared with the fed control (Fig 5b and 5e-h). The lipids can be comparably generated from both acetate and glutamine in feeding conditions, but mainly from glutamine in fasting conditions (Fig 5b and 5e-h). This result was further confirmed by ratiometric images that the ratio between glutamine derived lipid C–D signal (*I_2,182_*) and acetate derived lipid C–D signal (*I_2,135_*) was dramatically increased in the fasting mice (Fig 5i and 5j). Consistent with the metabolic imaging, RT-qPCR showed a significant increase in mRNA levels of *ACLY* and *GLD1* that are involved in glutamine mediated lipid synthesis (Fig 5k and 5l), but a marked reduction in mRNA levels of *ACSS2* and *ACACA* that are involved in acetate mediated lipid synthesis (Fig 5m and 5n) in the fasting liver. These data together suggest fasting drives substrate selection for glutamine to be preferentially incorporated into fatty acid synthesis. By contrast, acetate, which contributes less significantly to fatty acid synthesis, may become more utilized as an energy source during prolonged fasting.

Under normal feeding conditions, glycolysis and *de novo* lipogenesis were predominantly localized to hepatocytes near the central vein (CV) (Fig 5b), whereas gluconeogenesis and β-oxidation are enriched in periportal veins (PV) ^53^. It was corroborated by images showing a higher redox ratio (NADH/FAD) near the CV, while a lower redox ratio at the periportal region (Fig 5o and 5p). Strikingly, we observed that this zonal organization was reshaped upon fasting, with a notably transformed redox pattern (Fig 5o and 5p) and enhanced lipogenesis observed in periportal and intermediate hepatic zones (Fig 5b). Fasting induced zonation transformation was further validated by hyperspectral SRS imaging of C–H region and un-supervised K-means clustering analysis (Fig 5q and 5r). In control mice, the CV regions maintained a distinct spatial compartmentation of chemical clusters from that of PV, manifesting in clusters C3, C4, and C6 localized to the CV region (Fig 5q and 5r) and clusters C5 and C8 to the PV region (Fig 5q and 5r). However, in fasting mice, this spatial compartmentation became less defined. Notably, CV-specific clusters (C3, C4, and C6) were extended to the periportal region, indicating a convergence of chemical composition likely driven by lipid remodeling (Fig 5q). These relocated clusters were enriched in lipid signatures, as evidenced by their characteristic Raman spectral features with dominant peaks at 2850, 2880, and 3015 cm⁻¹ (Fig 5r), suggesting enhanced lipid synthesis as a potential driver of the altered metabolic zonation observed under fasting conditions. Together, these findings suggest that nutritional stress induces dynamic reprogramming of liver zonation, reflecting shifts in substrate utilization and spatial metabolic compartmentalization.

### SuMMIT-SRS unveils lipid and protein metabolic disorders in aging human induced neurons

In contrast to the remarkable metabolic flexibility of the liver in normal physiology, neurons exhibit limited metabolic plasticity and heightened vulnerability to metabolic stress ^54^. This is particularly evident in aging neurons, which undergo extensive metabolic rewiring as characterized by mitochondrial dysfunction, elevated reactive oxygen species (ROS) production, a shift toward aerobic glycolysis, impaired protein synthesis, and disrupted lipid metabolism ^55^. To visualize aging-associated alterations of protein and lipid metabolism in neurons, we applied SuMMIT-SRS to human induced neurons (i-Neurons). These neurons were directly transdifferentiated from fibroblasts (Fig 6a and 6b), which can retain donor-specific age-related epigenetic and metabolic signatures ^56–58^. We labeled young and old neurons (Fig 6c) with a combination of deuterium-labeled metabolic tracers (D-glucose, D-glutamine, D-methionine, D-tyrosine, and D-leucine) to compare their protein and lipid metabolic activities. After metabolic labeling, Raman microscopy revealed an increase of C–D intensity in old neurons (age 79, 82, 85) compared to young neurons (age 21, 22, 24) (Supplementary Fig 6a and Fig 6b). Quantitative analysis of CD/CH ratio indicated a higher macromolecular synthesis in old neurons (Supplementary Fig 6c). We further demonstrated that the increased C–D intensity was primarily driven by enhanced lipid synthesis (Supplementary Fig 6d and 6e). This increase coincided with notable rises in both the size and number of LDs in old neurons (Supplementary Fig 6f and 6g).

**Figure 6.**
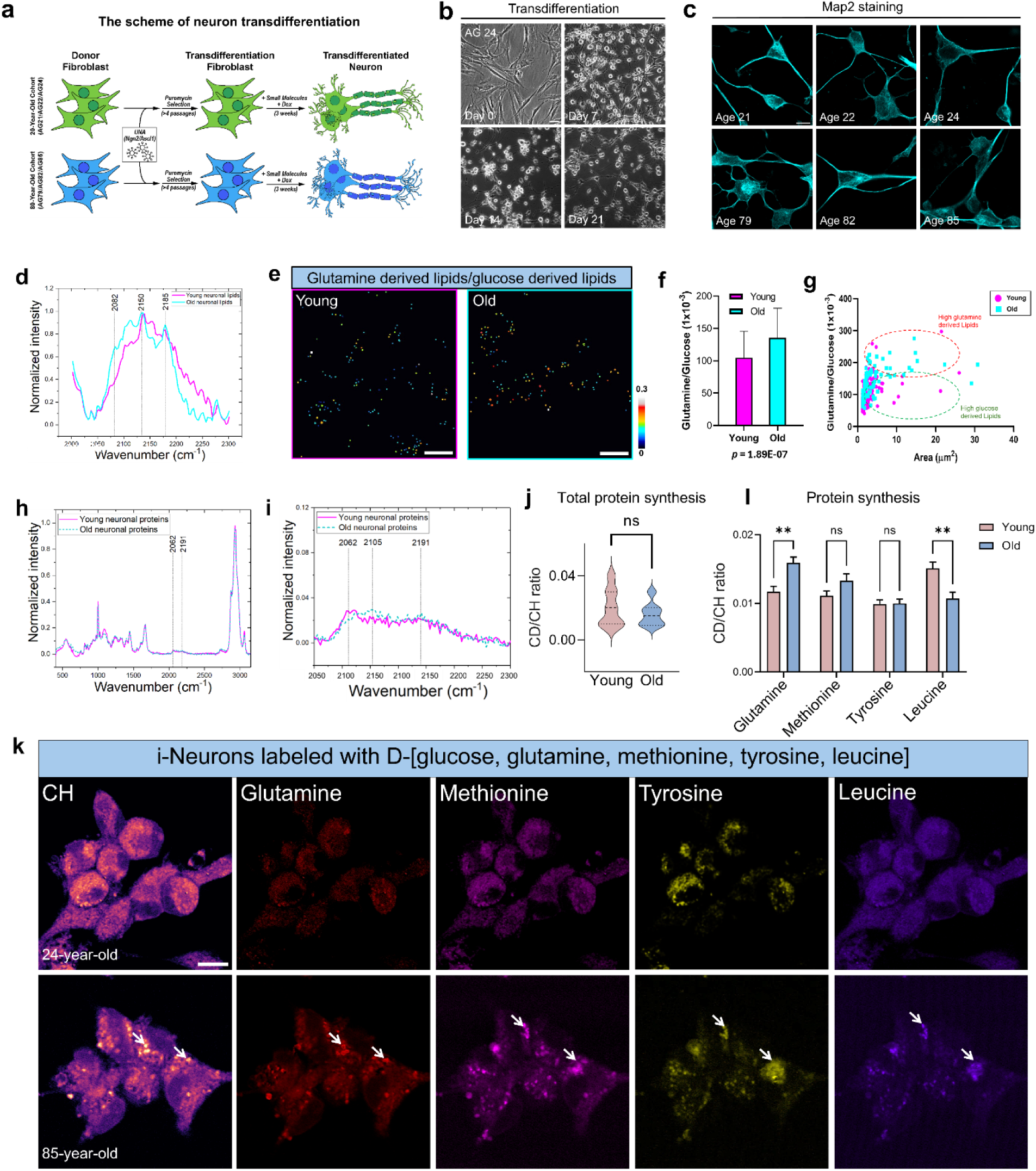
SuMMIT-SRS imaging unveils the metabolic pathway shift in aging i-Neurons. **(a)** Schematic depiction of direct conversion of donor fibroblasts to neurons. **(b)** Brightfield images showing the morphological characteristics of transdifferentiating neurons. **(c)** Map2 staining to confirm the identity of the converted neurons. **(d)** Raman spectra of the CD lipids from young and old neurons, respectively. Each spectrum was averaged from n=30 spectra collected from 3 independent groups. **(e-f)** A-PoD-processed high-resolution ratiometric images of glutamine/glucose lipid signal in single lipid droplets (LDs). The color of each LD represents the ratio intensity map of glutamine/glucose derived lipid ratio value (e). Quantification of the mean intensity of the glutamine/glucose-derived lipid ratio (f)**. (g)** Correlation between LD size distribution and glutamine/glucose-derived lipid ratio. The plot shows a clear separation between the ratios of Glutamine- to Glucose-derived lipids in two age groups, particularly for larger LDs. **(h)** Raman spectra of protein signals from young and old i-neurons after 5 days of co-labeling with D-glucose, D-glutamine, D-methionine, D-tyrosine and D-leucine. **(i)** Zoomed-in view of the C–D region from (h). **(j)** Quantification of protein synthesis by the ratio of CD (AUC) to CH (AUC) based on the Raman spectra in (i). n=15 spectra per group. **(k)** Images showing SuMMIT-SRS-detected single channel of glutamine-, methionine-, tyrosine-, and leucine-derived protein signals in the young and old neurons. Arrows indicate protein aggregations. Scale bar: 10 µm. **(l)** Quantification of protein synthesis from each deuterium labeled precursor by the CD/CH ratio based on the unmixed SRS signals in (K). n=15 ROIs per group. Values in (j, l) are mean ± SEM. ns, not significant; **, p < 0.01 by Student’s *t*-test.

SuMMIT-SRS detected a shift in the carbon source for lipogenesis in aged neurons: glutamine became the dominant contributor over glucose, as indicated by glutamine-to-glucose–derived lipid spectral shape (Fig 6d). High-resolution A-PoD–assisted analysis ^59^ of individual LDs further revealed elevated glutamine-to-glucose–derived lipid ratio and this metabolic shift was predominantly localized to larger LDs (Fig 6e-g). This shift may represent a compensatory response to reduced redox capacity in aging neurons, as glutamine metabolism can generate reductive equivalents beneficial for lipid biosynthesis.

In addition to lipid metabolism, we examined protein synthesis pathways. While Raman spectra indicated that the overall level of protein synthesis was comparable between young and old neurons (Fig 6h–j), aged neurons exhibited a significant increase in glutamine-derived protein and a concomitant decrease in leucine-derived protein (Fig 6k and 6l). Together with the enhanced glutamine-driven lipogenesis, these findings suggest that aged neurons display increased glutamine uptake and utilization for macromolecule biosynthesis. Notably, whereas newly synthesized protein signals were evenly distributed in young neurons, aged neurons displayed C–D protein signals in aggregated structures (Fig 6k). These observations indicate that aging neurons undergo substantial remodeling of protein synthesis pathways, characterized by altered substrate preference and increased propensity for newly synthesized proteins to form aggregates.

### SuMMIT-SRS parses spatially resolved cell type-specific metabolic pathway in developing brain

Building upon insights gained from models containing simple cell types, we extended SuMMIT-SRS to a more physiologically relevant and complex system, the developing *Drosophila* brain. The brain comprises diverse cell types, including neuroblasts, neurons, and glia, which arise from distinct neuroblast lineages (Fig. 7a). These major cell populations can be distinguished by their characteristic spatial localization and morphological features. The identity and fate of neuroblasts are governed by evolutionarily conserved cell-fate determinants and signaling pathways, which coordinate lineage specification and differentiation during brain development ^60,61^. Understanding how metabolic reprogramming intersects with these developmental programs is essential for deciphering how neural stem cells balance self-renewal, differentiation, and energy demands during brain development.

**Figure 7.**
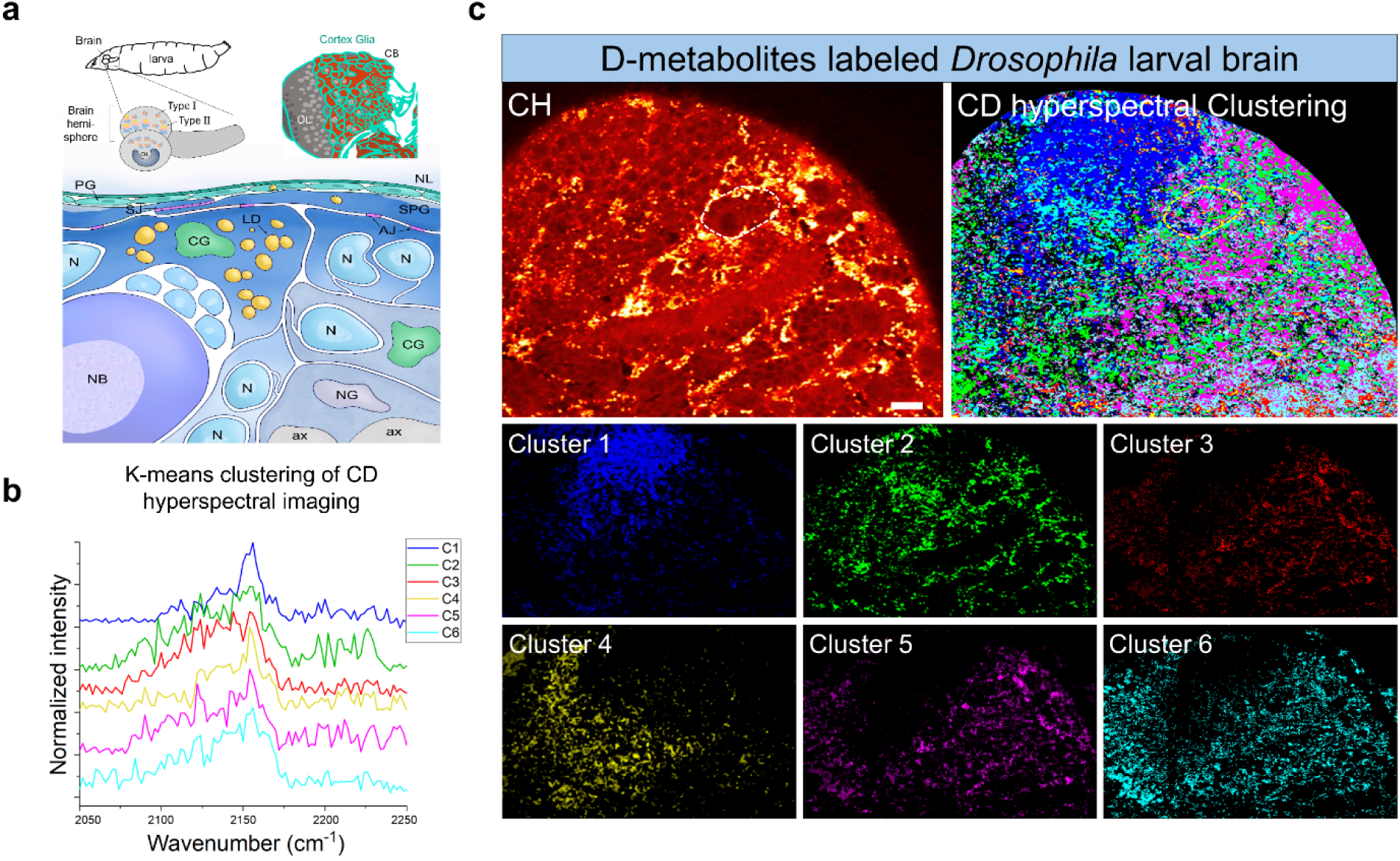
SuMMIT-SRS imaging uncovers cell-type-specific metabolic pathway in developing brain. **(a)** Cartoon depicting the anatomic structure of a *Drosophila* adult brain. CB: central brain, OL: optical lobe. The outermost layer of the brain is covered by the neural lamella (NL), an extracellular matrix that provides mechanical protection. Beneath the NL, perineurial glia (PG) form the surface layer, followed by subperineurial glia (SPG), which establish the blood–brain barrier through septate junctions (SJ) and adherens junctions (AJ). Within the brain cortex, cortex glia (CG) extend elaborate processes to enwrap neuronal cell bodies (N) and neuroblasts (NB), forming protective chambers and supporting neuronal metabolism. Cortex glia are also enriched in lipid droplets (LD), reflecting their role in energy storage and lipid homeostasis. Deeper within the neuropil, neuropil glia (NG) associate with neuronal processes and axons (ax), contributing to neurotransmitter clearance and metabolic support. Lipid droplets (LD) are mainly localized to glial cells, particularly cortex glia that enwrap neuronal stem cell niche. **(b-c)** Stimulated Raman Scattering Hyperspectral Imaging (SRS-HSI) combined with unsupervised K-means clustering applied to the C–D spectral region to analyze cell-type-specific metabolic activity. Each cluster, identified by a distinct color, corresponds to a unique chemical profile derived from the spectral data. The spatial localization of each color in the image reveals the distribution of these chemical features across the tissue. The spectral plot displays the average Raman spectra for each cluster, with the curve color matching the corresponding spatial region, thereby enabling direct correlation between chemical composition and tissue morphology or anatomic subregions. C, cluster. Each clustering image represents 10 brains from 3 independent experiments. A neuronal stem cell niche was highlighted in (c).

Using SuMMIT-SRS combined with 8 distinct deuterium-labeled metabolic tracers, we identified 6 main spectral clusters by K-means clustering of C–D hyperspectra (Fig 7b), which unveiled distinct cell type–specific metabolic profiles in *Drosophila* brain based on their unique spectral composition. Neurons in the optic lobe (OL), a region critical for visual information processing and synaptic maintenance, exhibited a unique metabolic signature (cluster 1) that was markedly different from neuroblasts (clusters 3, 5, and 6) and glial cells (cluster 2) (Fig 7b, 7c and Supplementary Fig 1a). Assisted by COMPASS-mediated spectral detection (Fig 4a), we found leucine-incorporated protein features (Raman peaks at 2,072, 2,133, and 2,188 cm⁻¹) were predominantly enriched in the neural stem cell niche in the central brain. This spatial pattern suggests that neuroblasts preferentially utilize leucine to fulfill their biosynthetic and proliferative demands while leucine metabolism may act as a critical link between nutrient availability and neurogenesis. In contrast, glial cells displayed dynamic lipid metabolism, with Raman spectral profile composed of lipids synthesized from glucose and fructose (Fig 7b, 7c and Supplementary Fig 1a). This suggests that glia is actively engaged in membrane biosynthesis and lipid-mediated signaling, both of which are essential for maintaining neural homeostasis and intercellular communication. Altogether, SuMMIT-SRS enables us to profile cell-type–specific metabolic pathway and spatially resolve cell-type–specific substrate preferences across all major brain cell populations, including glia, neuroblasts, and neurons.

## Discussion

Increasing evidence highlights that the divergent allocation of metabolites toward the synthesis of distinct macromolecular classes (such as nucleic acids, proteins, and lipids) reflects the functional state of the cell ^62,63^. We developed SuMMIT-SRS microscopy, a platform that integrates SRS microscopy with deuterium-labeled metabolic probes and linear unmixing algorithms as well as AI models to visualize and quantify the syntheses of major macromolecules, with high spatiotemporal resolution. SuMMIT-SRS enables simultaneously in situ tracing of multiple pathways of proteins, lipids, and nucleic acids biosynthesis, providing unprecedented insight into how distinct metabolic precursors are differentially allocated across macromolecular pools. We applied this platform to metabolic gene mutant flies and observed a striking increase in glucose- and glutamine-derived lipids, accompanied by reduced acetate-derived lipids. In tumor organoids, the analysis revealed that invasive networks are marked by elevated protein synthesis, diminished lipid synthesis, and differential routing of D-metabolite precursors into macromolecular production. In intermittently fasting mouse liver and aged human neurons, we found that glutamine supersedes glucose as the predominant substrate for lipid droplet biogenesis. Finally, studies in developing *Drosophila* brains uncovered region- and cell type–specific preferences in metabolic substrate utilization. Together, these findings highlight the versatility of SuMMIT-SRS imaging platform in uncovering context-dependent metabolic rewiring across species, tissues, and disease states. This imaging approach lays the foundation for super-multiplexed metabolic imaging, enabling simultaneous, pathway-resolved, and spatially precise visualization of diverse biosynthetic processes at subcellular resolution.

SuMMIT-SRS imaging offers high spatial resolution in a non-destructive manner. This capability addresses the growing need for in situ mapping of tissue metabolic heterogeneity, particularly in complex or spatially organized biological systems. SuMMIT-SRS imaging, when integrated with single-cell spatial omics and functional analyses in animal models, offers powerful potential for elucidating the metabolic architecture of complex and highly heterogeneous tissues. Moreover, it facilitates the exploration of cell type–specific metabolic programs and how they are shaped by genetic background, drug manipulation, developmental stage, or disease-associated mutations.

SuMMIT-SRS enables precise resolution of organelle-specific metabolic activity, particularly when combined with high-resolution modalities such as A-PoD-SRS. Although its spatial resolution and multiplexing capability may not reach the levels achieved by advanced super-resolution fluorescence imaging techniques, SuMMIT uniquely reveals metabolic pathway–dependent molecular diversity at the nanoscale. The integration of AI-driven analytical tools further enhances SuMMIT’s ability to detect subtle metabolic remodeling at the organelle level with high throughput across diverse biological contexts. Furthermore, by pairing SuMMIT with two-photon excitation fluorescence (TPEF) microscopy, which detects endogenous metabolic cofactors such as NAD(P)H and FAD to infer the redox state and metabolic activity, we can further delineate the whole picture of spatial architecture of cellular metabolism.

The SuMMIT labeling strategy can be readily expanded to a broader combination of other essential and non-essential amino acids, as well as other metabolic precursors, by incorporating diverse isotope labeling schemes. For instance, combining ^13^C or ^15^N isotopes with deuterium at defined atomic positions may result in distinguishable Raman spectral shifts, enabling the resolution of both atom origin and metabolic pathway engagement in a super-multiplexed way. Specifically, dual labeling with deuterium and ¹³C isotopes in acetate induces a characteristic red shift (∼10 cm⁻¹) in lipid Raman spectra while preserving the overall spectral shape (Supplementary Fig 1a), suggesting a distinctive chemical environment between the ¹³C–D and ¹²C–D bonds. This observation highlights the potential to utilize position-specific isotopologues as orthogonal probes for temporally and spatially resolved labeling of distinct acetate–derived lipid populations within the same sample, enabling applications such as lineage tracing and dynamic metabolic mapping.

Additionally, SuMMIT-SRS enables quantification of precursor-derived macromolecule turnover (half-lives) during aging and disease progression, providing key insights into the metabolic stability, remodeling capacity, and adaptability of cells in dynamic physiological contexts. Given that SRS microscopy is non-invasive, high resolution and quantitative, SuMMIT is particularly well-suited for live-cell pulse-chase experiments, enabling real-time tracking of labeled metabolite incorporation and degradation under controlled conditions. This dynamic imaging capability provides a powerful platform to investigate the temporal dimension of metabolic flux, offering a practical and informative means to address fundamental biological questions and deepen our understanding of both homeostatic and dysregulated cellular metabolism.

Our approach has inherent limitations related to labeling efficiency and detection sensitivity. It relies on deuterium incorporation as a proxy for metabolic flux, which can be influenced by variable precursor uptake and competition with unlabeled substrates. Low labeling efficiency may result in intracellular concentrations that fall below the Raman detection threshold. Moreover, metabolic transformations such as deamination or isotope exchange can lead to the irreversible loss of labeled atoms before incorporation into target macromolecules. While the AI-assisted COMPASS framework enhances spectral interpretation, it relies on high-quality reference libraries and is currently limited to known metabolite types, limiting its applicability in complex or poorly characterized systems.

## Methods

### SRS microscopy

We used an upright laser-scanning microscope (DIY multiphoton, Olympus) equipped with a 25× water objective (XLPlanN, WMP2, 1.05 numerical aperture, MP, Olympus), which was optimized for near-infrared throughput and transmission for SRS imaging. A picoEMERALD system (Applied Physics & Electronics) supplied a synchronized pulse pump beam (with tunable 720–990 nm wavelength, 5–6 ps pulse width and 80 MHz repetition rate) and Stokes beam (with fixed wavelength at 1031 nm, 6 ps pulse width and 80 MHz repetition rate). The Stokes beam was modulated at 20 MHz by an electronic optic modulator. Transmission of the forward-going pump and Stokes beams after passing through the samples was collected by a high numerical aperture (1.4) oil condenser. A high optical density bandpass filter (950 nm, Thorlabs) was used to block the Stokes beam completely and to transmit only the pump beam onto a large area Si photodiode for the detection of the stimulated Raman loss signal.The output current from the photodiode was terminated, filtered, and demodulated using a lock-in amplifier operating at 20 MHz. The demodulated signal was then fed into the FV-OSR software module (Olympus) integrated with the FV3000 microscope system to generate images during laser scanning. All acquired images were of size 512 × 512 pixels, with a dwell time of 80 μs and an imaging speed of approximately 23 seconds per frame. For multichannel SRS imaging, the pump wavelength (*λ*_pump_) was tuned so that the energy difference between pump and Stokes beams matched with the vibrational frequency. To minimize background interference, a background image was obtained at 1900 cm^-1^and subtracted from all SRS images using Fiji (ImageJ) software.

### SRS hyperspectral imaging

SRS-hyperspectral images were collected from the same setup described above. The hyperspectral imaging stacks of C–D region were taken with 100 spectral steps at 20 μs pixel dwell time, covering the region from 2,000 to 2,300 cm^−1^. The intensity profiles of the hyperspectral stacks from the regions of interest were plotted in ImageJ and further processed by baseline correction, smoothing and normalization in Originlab software (Origin Lab Corporation).

### Spontaneous Raman spectroscopy

Raman spectra of fixed cells or tissues were acquired using a confocal Raman microscope (XploRA PLUS, Horiba) connected to a Raman spectrometer. A diode line focus laser with a wavelength of 532 nm (∼40 mW at the sample) was utilized, and the laser beam was focused onto the cells using a 50× air objective (MPlanN, 0.75 numerical aperture, Olympus). The laser power was optimized to ensure cell integrity and prevent damage. Detection of the Raman signal was performed using a cooled charge coupled device (CCD) detector fitted to a spectrometer with a grating of 2400 grooves per mm. The acquisition time for each Raman spectrum was 60 seconds. The instrument calibration was validated using the silicon line at 520 cm^−1^. For cultured cells, the Raman background of water and cover glass is removed by subtracting the signal in cell-free areas from the signal collected in cells. For tissue with detectable autofluorescence background such as liver, the acquisition spot on the tissue was first photobleached by the laser for several minutes before spectra acquisition. All ratio calculations were conducted on the raw data before applying any normalization or baseline correction procedures. Data analysis and processing were performed using Originlab software. Data visualization was done in GraphPad.

### Cell culture, labelling and imaging

HeLa (ATCC) cells were maintained according to the instructions from ATCC in the DMEM medium. For labelling experiments, cells were seeded onto coverslips (No. 1, FisherBrand) and the culture medium was replaced with customed DMEM (ThermoFisher) media (Supplementary Table 2) supplemented with 10% fetal bovine serum (FBS) and individual or combined deuterium-labeled metabolites (Supplementary Table 1) at their standard concentration for the designated duration. Following labeling, the medium was replaced with PBS or cells were fixed with 4% PFA for 30 min, and the coverslip was sandwiched onto a glass slide (1 mm thick, VWR) with an imaging spacer (0.12 mm thick, SecureSeal) for imaging.

### Breast Cancer spheroids culture

MDA-MB-231 Triple Negative Breast Cancer cells were purchased from ATCC. MDA-MB-231 were cultured in high glucose Dulbecco’s modified Eagle’s medium (DMEM, Gibco), supplemented with 10% fetal bovine serum (FBS, Corning) and 0.1% gentamicin (Thermo Fisher Scientific). Cells were maintained at 37°C and 5% CO2 in a humidified environment.

### Preparation of collagen I gel for cancer organoids

To generate cell-laden Collagen I gels for Raman imaging, we adapted the approach of Leineweber et al. with minor adjustments ^47^. 12 mm glass coverslips were immersed in 100% ethanol for 1 hour, air-dried on lint-free wipes, and exposed to UV light for 20 min. The coverslips were then transferred to a 24-well plate and surface-activated in a plasma cleaner. Type I collagen gels were prepared by combining cells suspended in culture medium 1:1 with 10× reconstitution buffer before mixing with soluble rat tail type I collagen in acetic acid (Corning). The pH of the resulting mixture was adjusted to ∼7.0 by adding ∼2 µL of 1 M NaOH and 50 µL of the resulting mixture was dispensed onto each coverslip and allowed to polymerize at 37 °C for 30 minutes. Each gel had a final Collagen I concentration of 6 mg mL⁻¹ seeded with 6250 MDA-MB-231 Wild-Type cells. Standard medium was refreshed every other day for the first 4 days. After the 4 days, the gels were placed in glucose-, serum-, glutamine- and pyruvate-free DMEM (Gibco A1443001) for a 12h nutrient-depletion step, followed by 3 days in complete DMEM supplemented with D-metabolites and 10% FBS. Gels were rinsed in PBS, fixed for 20 min at room temperature with 4% paraformaldehyde, washed three times in PBS, and stored at 4°C until mounting on microscope slides.

### Macromolecule isolation from cultured cells

Hela cells were first cultured in 25 cm^2^ culture flask in complete medium to 50% confluence. The medium was then replaced with DMEM substituted with D-metabolites. The medium was replaced with fresh medium every 2 d. After 3 days, cells were dissociated and collected, and DNA, RNA and proteins were extracted using TRIzol Reagent (ThermoFisher) according to the manufacturer’s instructions. For lipid extraction, cells were collected and suspended in 1 ml PBS and mixed with 1.3 ml chloroform and 2.7 ml methanol. The solution was centrifuged at 4,000 r.p.m. for 5 min. The supernatant was transferred to a new clean tube and mixed with 1 ml of 50 mM citric acid (Sigma-Aldrich), 2 ml distilled water and 1 ml chloroform. After briefly shaking and mixing, the tube was centrifuged at 4,000 r.p.m. for 10 min. The material separated into three phases: a water–methanol upper phase, a middle phase of precipitated protein and a lower chloroform phase. Lipids were dissolved in the lower chloroform phase. The chloroform phase was separated, dropped on a cover glass and air-dried.

### Drosophila experiment

Wild-type flies (w1118 stock #5905) were initially acquired from the Bloomington Stock Center (DBSC) and have been maintained in the laboratory for multiple generations. The flies were kept under standard conditions, including a temperature of 25 °C, a humidity level of 60%, and a 12-hour dark/light cycle. They were fed with corn meal-based fly food (Nutri-Fly, Cat. No. 66-113; Genesee Scientific Corporation), following established protocols. For D-metabolite labeling experiments, individual or combined deuterium-labeled metabolites were substituted for the corresponding components in the home-made food and administered for the designated period. The food recipe was prepared as previously described (Supplementary Table 3) ^49^.

### Mouse labelling

All mouse experiments were conducted in adherence to the experimental protocol approved by the Institutional Animal Care and Use Committee (AC-AAAQ0496). C57BL/6J mice (The Jackson Laboratory, stock no. 000664) were used for all experiments. For normal conditions, mice were given drinking water containing D-labels at the designed concentration. Mice had free access to regular chows during this time. For intermittent fasting, the experimental group of mice was subjected to 2:1 (2 days of feeding, 1 day of fasting) pattern intermittent fasting, during which they were provided with access to drinking water containing the same concentration of D-labeled metabolites as the control group. During feeding phases, the mice were given normal chow diet. The entire treatment lasted for 10 days (Supplementary Fig 5a).

### Tissue slicing

After treatment and labelling, the mice were anaesthetized, killed by cervical dislocation and organs were collected. The tissues were fixed with PFA for more than 48 h after collection. Tissues were then embedded in 4% agarose gel (Sigma-Aldrich) and sliced into 120 μm thin slices using a vibratome (Leica). Brain was sliced coronally. Random orientation was used for other tissues. The tissue slices were collected and sealed between a glass slide and coverslip.

### Tissue processing

To remove lipid, tissue slices were soaked in pure methanol for 48 h and then washed with PBS. To extract lipid from adipose tissue, 1 g of tissues were collected, ground and suspended in 1 ml PBS, and then mixed with 2.6 ml chloroform and 5.4 ml methanol. The suspension was centrifuged at 4,000 r.p.m. for 5 min. The supernatant was transferred to a new clean tube and mixed with 2 ml of 50 mM citric acid (Sigma-Aldrich), 4 ml water and 2 ml chloroform. After brief shaking and mixing, the tube was centrifuged at 4,000 r.p.m. for 10 min to separate the phases. The lower chloroform phase containing lipids was collected and solvent was evaporated to obtain tissue lipid extract.

### Fibroblast-to-neuron transdifferentiation

#### Lentiviral Production

Lenti-X 293T (Takara #632180) cells were cultured in DMEM (high glucose, Life #11965118) supplemented with 10% (v/v) fetal bovine serum (Life #26140079) at 37 °C with 5% CO2. For lentiviral production, the Lenti-X 293T cells were passaged to a fresh plate at ∼50% confluency. Within 24 h of a fresh passage, the newly-seeded cells were co-transfected with psPAX2 (Addgene #12260), pMD2.G (Addgene #12259), and pLVX-UbC-rtTA-Ngn2:2A:Ascl1 (UNA; Addgene #127289) at a 1:1:2 mass ratio using the JetPEI transfection reagent (VWR #89129-916). The supernatant fraction of the transfected cells was collected 48 h, 72 h, and 96 h and pooled at 4 °C. The collected virus was added to Lenti-X concentrator (Takara #631232), incubated at 4 °C for 30 min, and centrifuged at 1500 x g for 45 min at 4 °C. Following centrifugation, the lentivirus-containing pellet was resuspended in 1X dPBS (Life #13190250), the viral titer was measured using Lenti Go-Stix (Takara #631280), and the purified virus was diluted to the desired titer for storage at -80 °C.

#### Transdifferentiation

In brief, fibroblasts (Supplementary Table 4) were maintained in TFM (DMEM, high glucose; 15% (v/v) fetal bovine serum; 1X MEM NEAA, Thermo #11140050) at 37 °C with 5% CO2. TFM was replenished every 2-3 days, and fibroblasts were always maintained at >30% confluency. For passaging, fibroblasts were detached with TrypLE Express (Life #12604013) for 5 min and split at 1:2 or 1:3 split ratios. Long-term fibroblast stocks were resuspended in BamBanker (Bulldog Bio #BB05) for storage in liquid nitrogen. To generate transdifferentiation-competent fibroblasts, 80% confluent cells were seeded on 6-well plates (Fisher #FB012927) incubated in 500 μL TFM with lentivirus containing the UNA expression vector in 5 μg/mL polybrene (Sigma #TR-1003-G) for 8 h at 37 °C. Afterwards, the transduction volume was increased to 2 mL for an additional 36 h incubation. The transduced fibroblasts were then incubated in TFM-P (TFM with 1 μg/mL puromycin, Thermo #A1113803) for at least 4 passages to select for transdifferentiation-competent fibroblasts. Cultures with death rates greater than 50% after selection were discarded.

Plates for transdifferentiation were coated with 10 μg/mL poly-D-lysine (PDL; Sigma #P6407-5MG) and 0.0001% (w/v) poly-L-ornithine (PLO; Sigma #P4957-50ML) for 24 h and 10 μg/mL laminin (Sigma #L2020-1MG) for an additional 24 h. The coated plates were seeded with 300% confluent UNA fibroblasts. Transdifferentiating neurons were cultured for three weeks in NK media (NK) (0.5X DMEM/F-12, GlutaMAX supplement, Life #10565042; 0.5X Neurobasal, Life #21103049; 1X B-27 supplement, Life #17504044; 1X N-2 supplement, Life #17502048; 20 μg/mL laminin; 400 μg/mL dibutyryl-cAMP (db-cAMP), SelleckChem #S7858; 2 μg/L doxycycline, Sigma #D9881-10G; 5 μM dorsomorphine homolog 1 (DMH1), Fisher #412610; 0.5 μM LDN-193189, Fisher #50176042; 0.5 μM A83-01, Stem Cell Technologies #72022; 5 μM forskolin, Stem Cell Technologies #72114; 3 μM CHIR-99021, Fisher #442310; 10 μM SB-431542, Fisher #161410; 10 U/mL penicillin-streptomycin, Thermo #15140122) with media changes every 48-72 h. Mature neurons were passaged to laminin-coated coverslips (Fisher Scientific #NC0937501) for Raman spectroscopy or PDL/PLO/laminin-treated Ibidi chambers (Ibidi #81811) for immunofluorescence and maintained in BrainPhys+GF media (BP-GF) (1X BrainPhys, Stem Cell Technologies #05790; 1X B-27 supplement; 1X N-2 supplement; 400 μg/mL db-cAMP; 1 μg/mL laminin; 20 ng/mL BDNF, PeproTech #450-02-50UG; and 20 ng/mL GDNF, PeproTech #450-10-50UG) for 1 week until imaging. Transdifferentiation progress was monitored using a Zeiss Axio-Vert.A1, which was also used to acquire brightfield images with a 20x objective lens.

#### Immunofluorescence

Neurons cultured in Ibidi chambers were fixed with 4% paraformaldehyde (Fisher #50980492) with gentle rocking at 4 °C for 45 min, washed three times with 1X dPBS, permeabilized with 1X dPBST (1X dPBS supplemented with 0.1% (v/v) Triton X-100, Sigma #X100-500ML) with 5% (v/v) donkey serum (Jackson ImmunoResearch #017-000-121) for 45 min at 25 °C, washed three times with 1X dPBST, incubated with Chicken Anti-Map2 primary antibody (Invitrogen PA1-10005, 1:250 dilution) in 1X dPBST with 0.1% (w/v) bovine serum albumin (Fisher #10773877) for 16 h at 4 °C, washed an additional three times with 1X dPBST, and incubated with Donkey Anti-Chicken IgY, Alexa Fluor 488 (Thermo #A-78948, 1:2000 dilution) for 2 h at 25 °C. The stained cells were mounted in Prolong Diamond Antifade mountant (Thermo #P36971), cured for 24 h at 25 °C, and stored at 4 °C until imaging. Confocal imaging was performed on a Zeiss LSM-880 confocal microscope with a 63x objective lens at the Salk Advanced Biophotonics Core. Images were processed using ImageJ and analyzed for Map2+ staining with the image thresholding function (n=5 replicates).

### AI for hyperspectral SRS imaging detection

#### Data Preprocessing and Augmentation

To bridge the spectral discrepancies between high-quality reference spontaneous Raman spectra and lower signal-to-noise ratio (SNR) pixel-level HSI spectra, we implemented specialized data augmentation techniques. These strategies simulate common spectral variations encountered in HSI, including noise, instrument shifts, and background interference, thereby generating training data more representative of real-world pixel spectra. Let S_ref_(ν)denote the reference Raman spectral intensity of a metabolite at wavenumber ν. Augmented spectra, 𝑆𝑆_aug_(𝜈), were generated using the following strategies:

To simulate shot noise and detector noise inherent in HSI, Gaussian noise was added to the reference spectra:

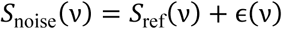

where 𝜖(𝜈) represents random noise sampled from a Gaussian distribution 𝑁(0, σ^2^). The standard deviation 𝜎𝜎 was varied, based randomly selected from a predefined range, to mimic diverse SNR conditions.

To account for potential instrument drift or calibration inaccuracies, the wavenumber axis of the reference spectra was randomly shifted:

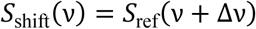

where Δ𝜈𝜈 is a shift value randomly sampled from a uniform distribution with δ defining the maximum shift range. HSI pixel spectra often contain background contributions from the sample matrix or solvent (e.g., water or PBS). This was simulated by adding authentic background spectra, such as a water spectrum 𝐵_water_(𝜈) extracted from HSI data to the reference spectra with a random scaling factor:

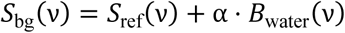

where 𝐵_water_(𝜈) is the Raman spectrum of water, and 𝛼 is a random scaling factor controlling the background contribution level.

These augmentation strategies were often applied concurrently. A comprehensively augmented spectrum S₍aug₎(ν) can be represented as:

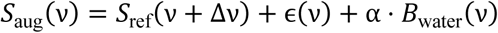

Applying these augmentations to each reference spectrum significantly expanded the training dataset, enabling the model to generalize across the spectral variability observed in HSI data.

#### Model Training

A one-dimensional residual neural network (1D-ResNet) architecture, featuring four residual blocks, was employed for feature learning and classification. The training proceeded in two stages:

Pre-training: The 1D-ResNet model was initially trained using the augmented reference spectral dataset. The objective was to establish baseline capabilities for distinguishing fundamental spectral features of different metabolites. Optimization was typically performed using the cross-entropy loss function:

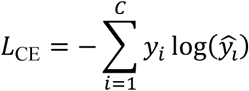

where *C* is the number of metabolite classes, *y_i_* is the ground-truth label (one-hot encoded), and ̂*y_t_* is the model’s predicted probability for class i.

Post-training / Fine-tuning: To enhance performance on real HSI data application and mitigate interference from tissue background signals, a crucial post-training phase was conducted. During this stage, the pre-trained model was fine-tuned using background spectra extracted directly from actual HSI images (potentially containing autofluorescence, water signals, etc.). These spectra were assigned to a distinct “background” class. This step significantly improved the model’s ability to differentiate target metabolite signals from complex real-world backgrounds, particularly within the low-SNR C-D stretching region.

#### Hyperspectral SRS Image Inference

The trained model was subsequently applied to new HSI datasets for pixel-wise inference.

For each pixel (𝑥, 𝑦) in the HSI dataset, its corresponding spectrum 𝑆_pixel_(𝑥, 𝑦, ν) was fed into the trained 1D-ResNet model. The model outputs the probability 𝑃 (𝑐 | 𝑆_pixel_(𝑥, 𝑦, ν)) for the pixel belonging to each metabolite class *c* and the background class.

To enhance prediction reliability and minimize false positives, an optimized probability threshold 𝑇_𝑐_ was applied for each class *c*. A pixel was assigned to class *c* only if its predicted probability 𝑃(𝑐 | 𝑆_pixel_) exceeded 𝑇_𝑐_ and was the maximum probability among all classes. Thresholds 𝑇_𝑐_ were typically determined as 0.75. The final classification results for all pixels were assembled to generate spatial distribution maps for each metabolite.

#### Model Interpretability

To understand the basis of the model’s predictions, Gradient-weighted Class Activation Mapping (Grad-CAM) was employed. Grad-CAM visualizes the wavenumber regions that were most influential in classifying a specific metabolite, indicating which parts of the spectrum the model primarily “focused” on. This provides intuitive interpretability for the model’s outputs and aids in verifying that the model learned relevant spectral features.

#### Quantitative RT-PCR

Total RNA from mouse livers was extracted by using RNeasy Mini Kit (QIAGEN, 74104) following manufacturer’s protocol. The extracted RNA was subjected to DNase I treatment to eliminate genomic DNA contamination. 500 ng of RNA was reverse transcribed to cDNA using iScript Reverse Transcription Supermix (Bio-Rad, 1708841). qRT-PCR was performed using Maxima SYBR Green qPCR Master Mix (Thermo Fisher, K0253) in the CFX96 Real-Time PCR Detection System (Bio-Rad). The expression levels of genes of interest were normalized to that of housekeeping gene (ACTB), and relative expression fold change was calculated using the 2^−ΔΔCt^ formula.

#### Optical redox imaging

The two-photon fluorescent microscopy was used to collect the NADH and FAD signals from the tissue. The NADH signal was excited by 780 nm, the emission wavelength was collected at 460 nm. The FAD signal was excited by 860 nm, the emission wavelength was collected at 515 nm. The ratiometric images were generated by FAD/NADH. The ratio was further quantified in ImageJ and plotted in GraphPad 10.

## Acknowledgements

We thank Bloomington Drosophila Stock Center, DSHB. We thank A.J. Z. for donating Plin1 mutant fly line. We acknowledge fundings from NIH R01GM149976, NIH U01AI167892, NIH 5R01NS111039, NIH R21NS125395, NIHU54 HL165443, NIH U54 DK 134301, NIH R01 HL170107, Sloan research fellow award, CZI Scialog award, and UCSD Startup funds to L.S., as well as NIH U54AG076043 (to R.F. and L.S.) and U54AG079759 (to R.F.). K.R. was funded by the Milton Safenowitz Postdoctoral Fellowship via the ALS Association (23-PDF-639). G.W.Y was funded by Grant 2023-332369 from the Chan Zuckerberg Initiative DAF, an advised fund of Silicon Valley Community Foundation, and the NIH (R01-HG004659 and R01-NS103172). S.I.F. and S.K.R. were supported by American Cancer Society RSG-21-033-01-CSM, National Cancer Institute CA274502, and a Prebys Research Heroes Award.

## Author contributions

L. S. conceived the idea of the project. Y. L. and L. S. designed the experiments. Z. L. developed the COMPASS spectral detection method. Y. L., Y. L., K.R. and S. R. performed the experiments and analyzed the data with the help from Z. L., S. Q., H. J., J. V., Z. B., S. F., G. Y., R. F., and L. S.. Y.L. wrote the first draft. Y.L. and L.S. finalized the manuscript with the input from all authors.

## Supplementary Figures

**Supplementary Figure 1.**
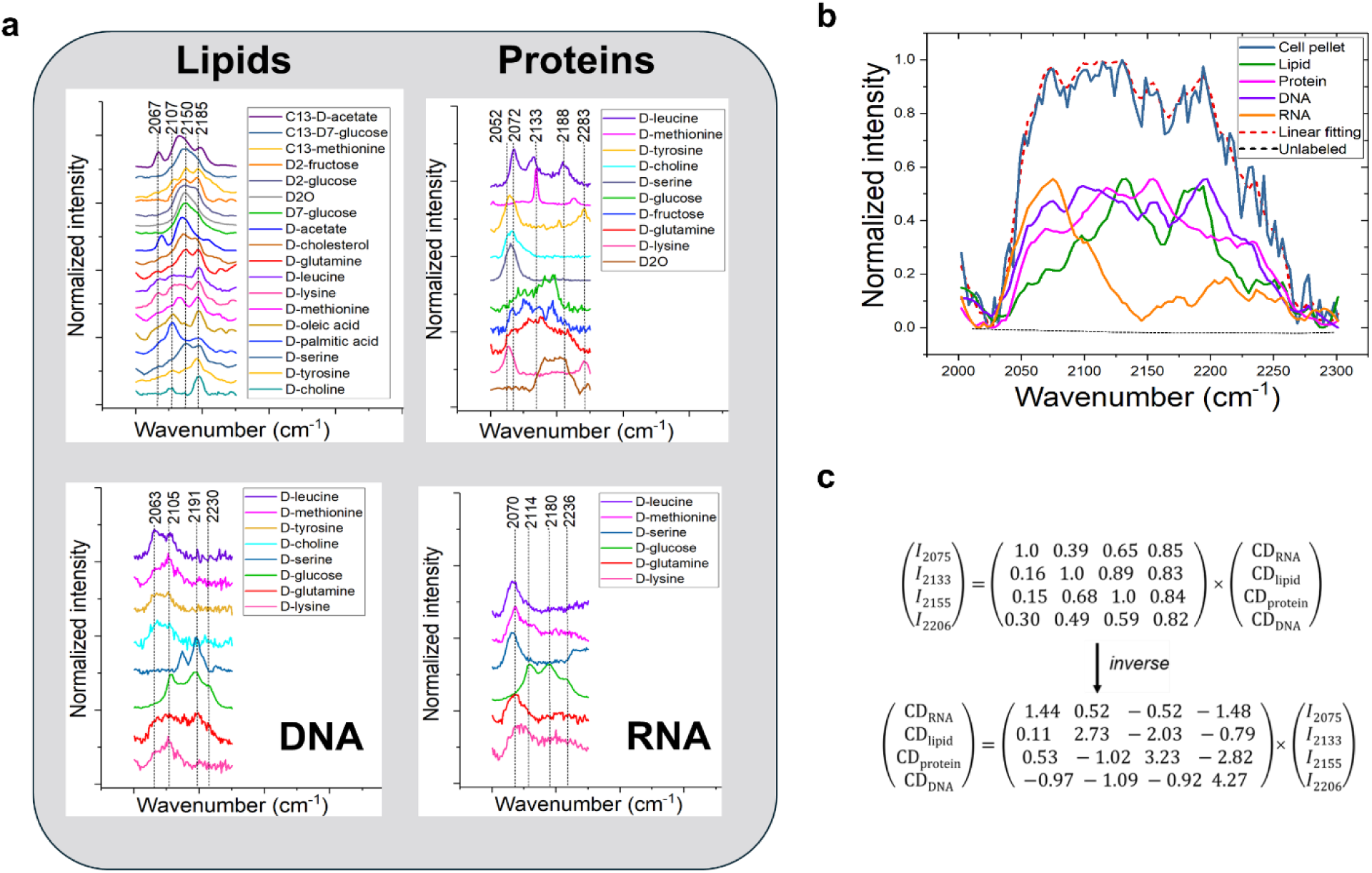
Spectral library of SuMMIT-SRS. **(a)** Raman spectra of D-metabolites labeled protein, lipid, DNA, and RNA spectra from Hela cells and fly tissues measured by spontaneous Raman microscopy. **(b)** Raman spectra of extracted proteins, lipids, DNA, RNA and overall cell pellets from D₅-glutamine–labeled Hela cell after culturing in medium supplemented with 4 mM D₅-glutamine for 72 h. The computerized Raman spectrum generated from linear fitting of standard protein, lipid, DNA, and RNA spectrum agree well with the spectrum from cell pellets. Each spectrum was averaged from n=3 independent experiments. **(c)** The factors used for spatial SRS signal linear decomposition algorithm for Fig 1c.

**Supplementary Figure 2.**
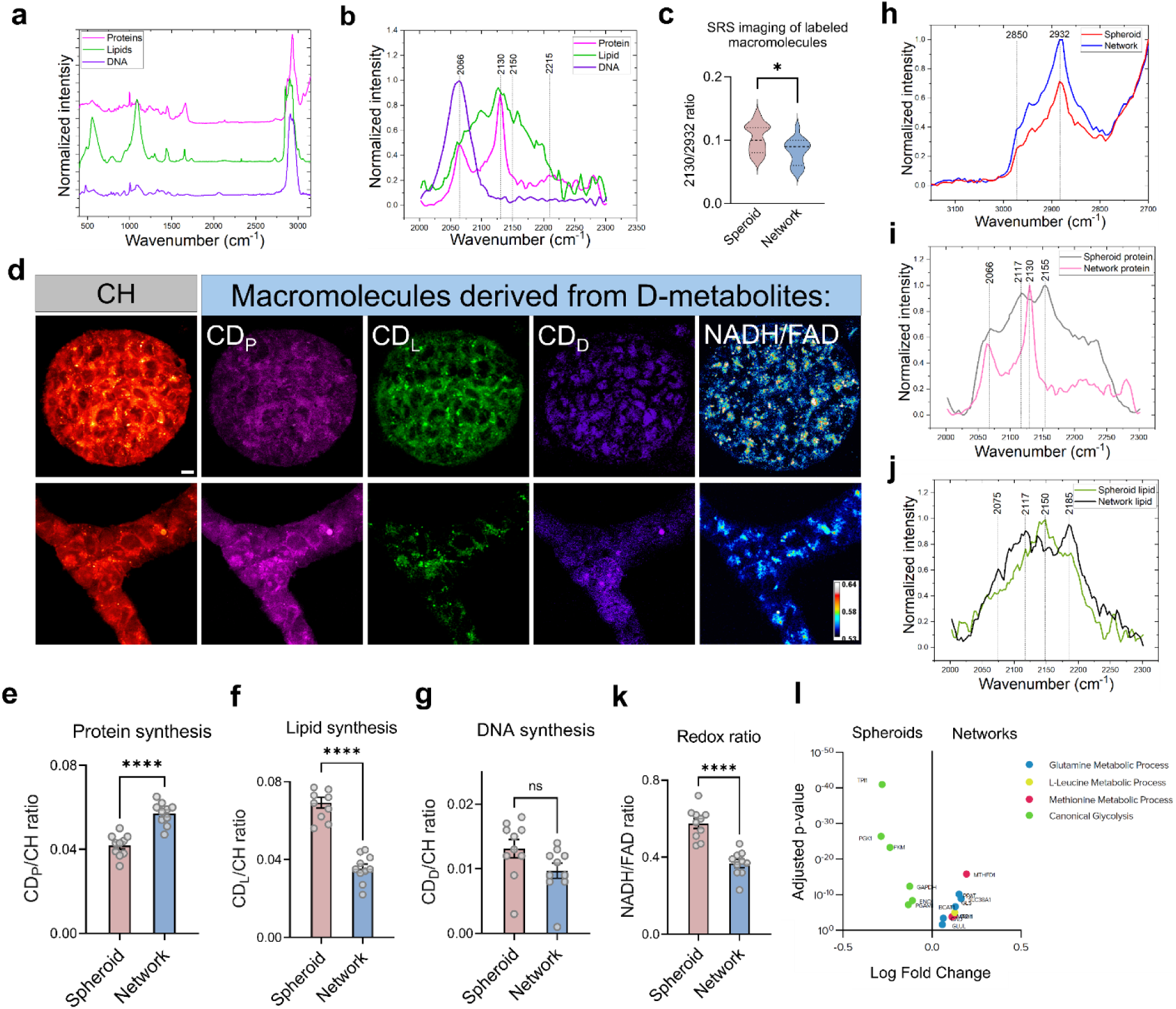
SuMMIT-SRS imaging examined the heterogeneity of breast cancer (BRCA) organoids. **(a)** Raman spectra of extracted proteins, lipids, and DNA from BRCA organoids with combinational labeling of D-glutamine, D-glucose, D-methionine, D-tyrosine and D-leucine measured by spontaneous Raman microscopy. Each spectrum was averaged from n=15 ROIs per group. **(b)** Zoomed-in CD Raman spectra of BRCA organoids with mixed labeling of D-glutamine, D-glucose, D-methionine, D-tyrosine and D-leucine. Each spectrum was averaged from n=15 ROIs per group. **(c)** Quantification of total macromolecule synthesis of spheroids and networks by the peak intensity ratio of 2130/2932 measured from SRS imaging. n=10 ROIs per group. **(d)** SRS imaging of total proteins (CH3), newly synthesized deuterium labeled macromolecules including proteins (CDP), lipids (CDL), and DNA (CDD). And two photon excited fluorescence imaging of redox ratio indicated by (NADH/FAD). Scale bar: 10 µm. **(e-g)** Quantification of protein (e), lipid (f), and DNA (g) synthesis as well as redox ratio (I) of tumor spheroids and networks with combination co-labeling of D-glutamine, D-glucose, D-methionine, D-tyrosine and D-leucine. **(h)** Raman spectra of pure protein signal of spheroids and networks treated with methanol wash. n=15 ROIs per group. **(i)** Raman spectra of pure CD protein signal of spheroids and networks treated with methanol wash. n=15 ROIs per group. **(j)** Raman spectra of pure CD lipid signal of spheroids and networks treated with proteinase K. n=15 ROIs per group. **(k)** Redox ratio imaging of spheroids and networks. n=10 ROIs per group. **(l)** RNA seq data analysis showing the upregulated metabolic pathways in networks. Values in (c, e-k) are mean ± SEM. ns, not significant; *, p < 0.05; ****, p < 0.0001 by Student’s *t*-test.

**Supplementary Figure 3.**
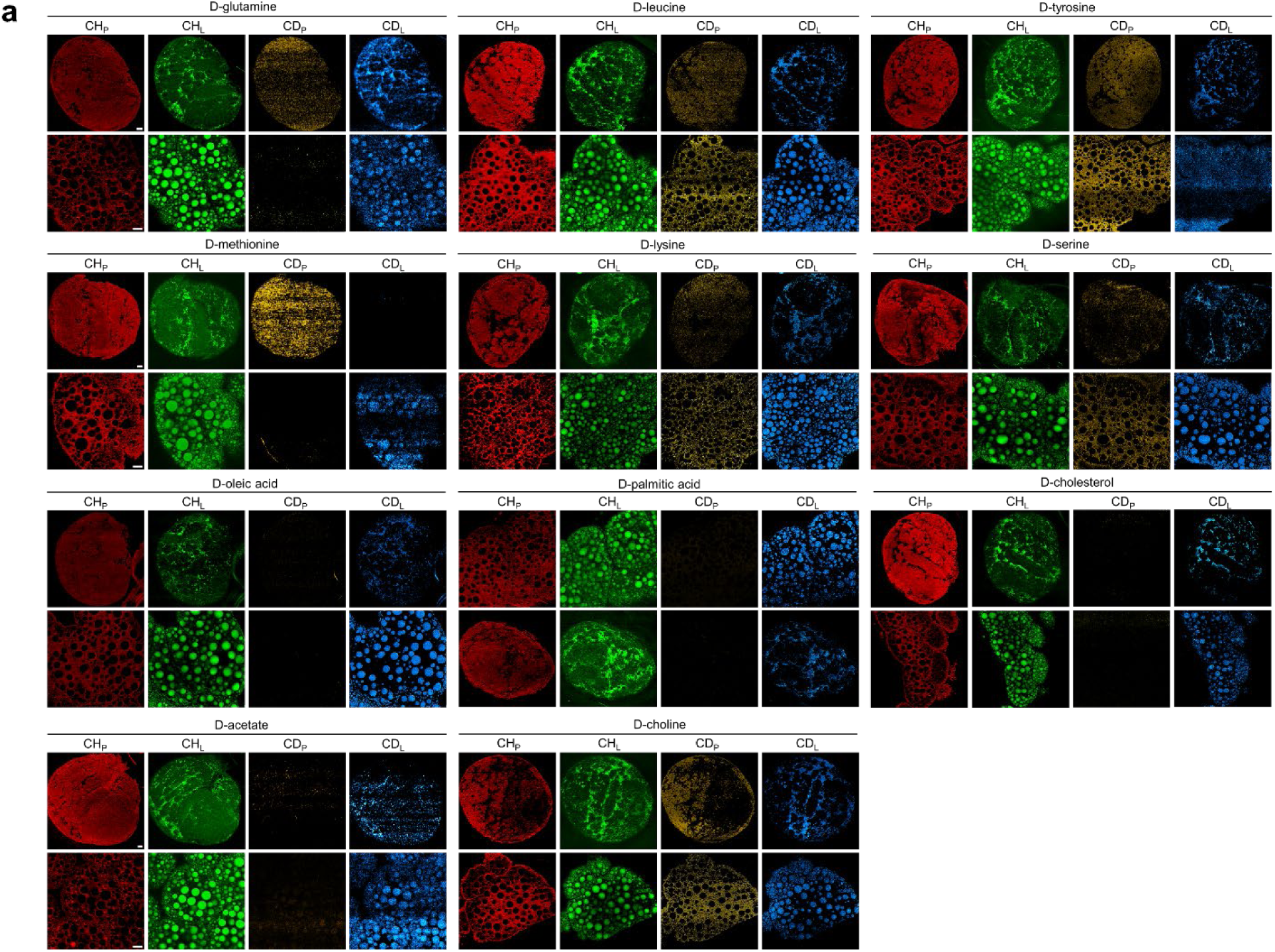
SuMMIT-SRS imaging for assessing macromolecular synthesis in *Drosophila* larvae. **(a)** As shown in the Fig 3a schematic: The parent flies were cultured in standard cornmeal foods without D-metabolites. The eggs were collected and subjected to standard cornmeal foods for 24 h development. And then the first instar larvae (24 h after egg lay) were transferred to the chemically synthesized food with D-metabolites for 3 days, after which the mid third instar larvae were sacrificed, and brains and fat bodies were dissected and subjected to SRS imaging. Following SRS imaging, the protein and lipid distribution were obtained by linear unmixing with the factors determined by the standard protein and lipid spectra measured from *Drosophila* brains and fat bodies, respectively. Each image represents 10 ROIs from at least 3 independent experiments.

**Supplementary Figure 4.**
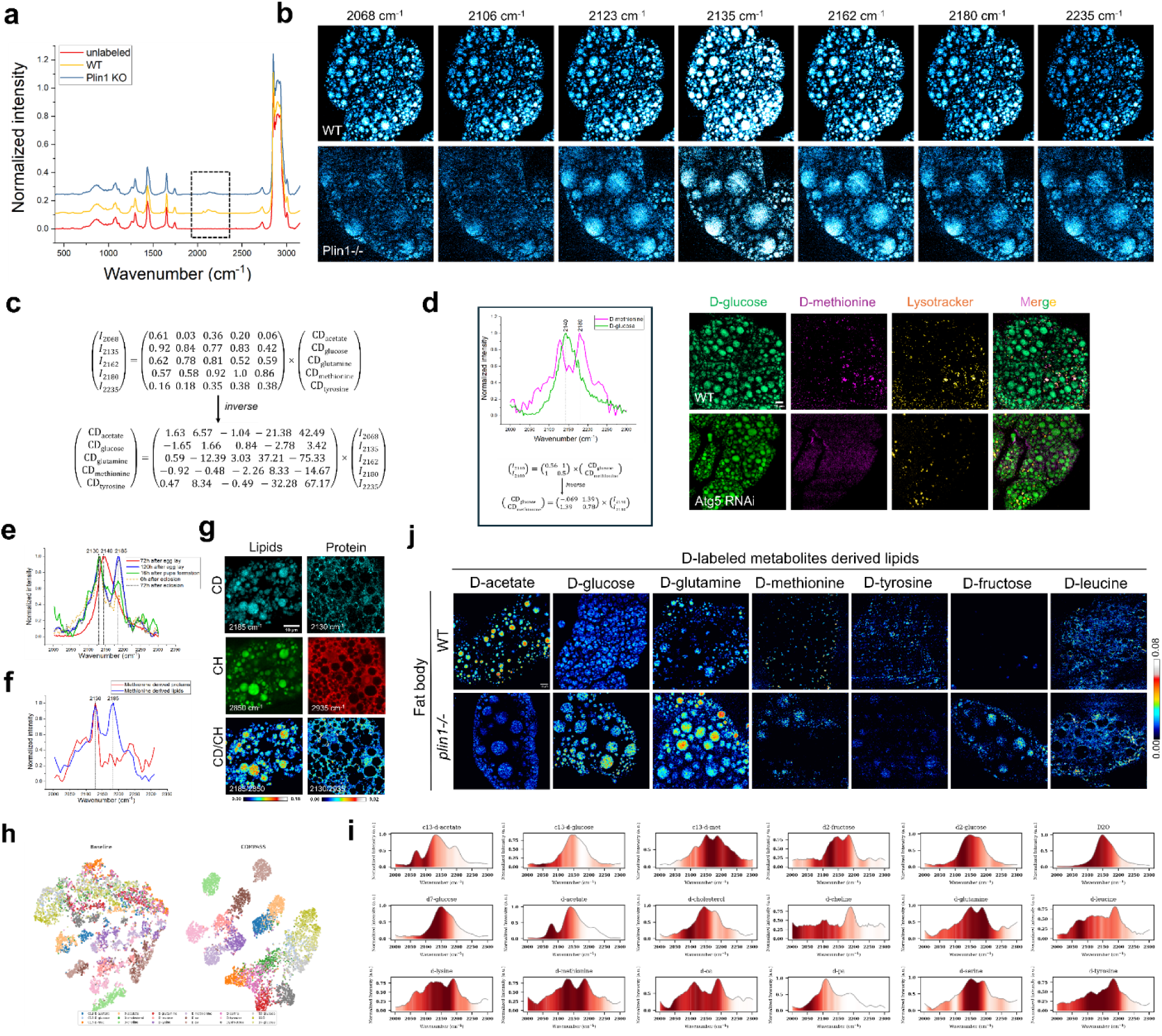
SuMMIT-SRS imaging to dissect metabolic reprogramming in *Drosophila* fat body. **(a)** Raman spectra of WT and Plin1 mutant fat bodies with a combinational labeling of D-acetate, D-glucose, D-glutamine, D-methionine, and D-tyrosine. **(b)** SRS images of WT and Plin1 mutant fat bodies collected at different wavelengths within the cell silence region before performing spectral decomposition. Scale bar: 10 µm. **(c)** Factors used for linear unmixing were determined by the standard spectra of each single labeling collected from fat bodies. **(d)** SRS imaging of D-glucose and D-methionine labeling in *Drosophila* third instar larvae. Notably, the D-methionine signal was found in the lysosome which was colocalized with lysotracker. And the signal disappeared upon *Atg5* knocking down. Scale bar: 10 µm. **(e)** Raman spectra of D-methionine labeled fat body at different developmental stages. **(f)** Raman spectra of D-methionine derived proteins and lipids extracted from adult fat bodies. **(g)** SRS images of D-methionine derived pure lipids and proteins at 2185 and 2130 cm^-1^, respectively. The ratiometric images indicating lipid and protein turnover were generated by the intensity ratio between CD-lipids (2185 cm^-1^)/ CH-lipids (2850 cm^-1^) and CD-proteins (2130 cm^-1^)/ CH-proteins (2935 cm^-1^). Scale bar: 10 µm. **(h)** UMAP visualization of C– D spectral data from 18 isotope-labeled metabolites under baseline analysis (left panel) compared with the COMPASS framework (right panel). Whereas baseline analysis shows substantial overlap among different metabolites, COMPASS achieves improved clustering and separation across metabolite classes. **(i)** Normalized Raman spectra (2,000–2,300 cm⁻¹) of each metabolite, highlighting the characteristic C– D vibrational signatures used for feature extraction and classification. The shaded regions indicate feature-weighted contributions learned by COMPASS, which enable accurate discrimination across metabolites. **(j)** SRS CD/CH ratiometric images of WT and Plin1 mutant fly fat bodies labeled with the indicated D-metabolite individually. Scale bar: 10 µm.

**Supplementary Figure 5.**
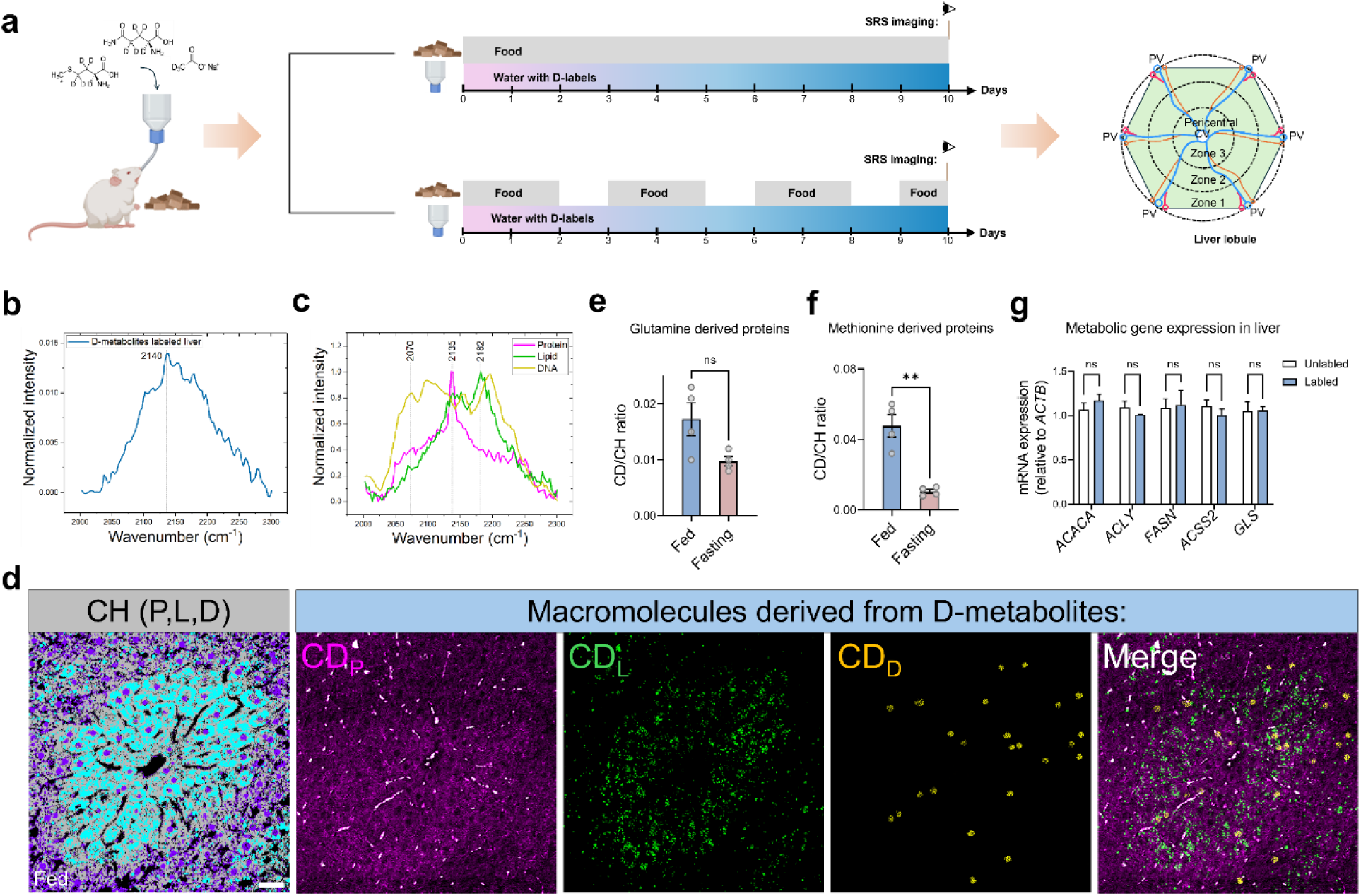
SuMMIT-SRS imaging of metabolic pathway remodeling in mouse liver under intermittent fasting. **(a)** Schematic of SuMMIT-SRS imaging timeline for assessing macromolecular synthesis in mice under different diet regimens. The control group was fed with the normal diet and drinking water with the D-metabolites for continue 10 days while the experimental group was subjected to 2:1 patten interm fasting. After labeling, the mice were sacrificed, and the liver was dissected out and sectioned to be examined with SRS imaging. **(b)** Raman spectra of overall C–D signal raised from the D-metabolites labeling in mouse livers under normal diet labeled with a combinatorial labeling with D-glutamine, D-methionine and D-acetate. Each spectrum was averaged from n=10 ROIs per group. **(c)** Raman spectra of CD-proteins, lipids and DNA extracted from D-metabolites labeled mouse livers under normal diet. Each spectrum was averaged from n=10 ROIs per group. **(d)** SRS imaging of C–H and C–D protein, lipid and DNA distribution of liver labeled with D-metabolites under normal diet. Each image represents 10 ROIs from at least 3 independent experiments. **(e)** Quantification of glutamine derived proteins in fed and fasting mice. n=15 ROIs per group. **(f)** Quantification of methionine derived proteins in fed and fasting mice. n=15 ROIs per group. **(g)** Quantification of the expression levels of the metabolic genes (*ACACA*, *ACLY*, *FASN*, *ACSS2*, *GLS*), which are involved in glutamine and acetate metabolic pathways. n=15 ROIs per group. Values in (e-g) are mean ± SEM. ns, not significant; **, p < 0.01 by Student’s *t*-test.

**Supplementary Figure 6.**
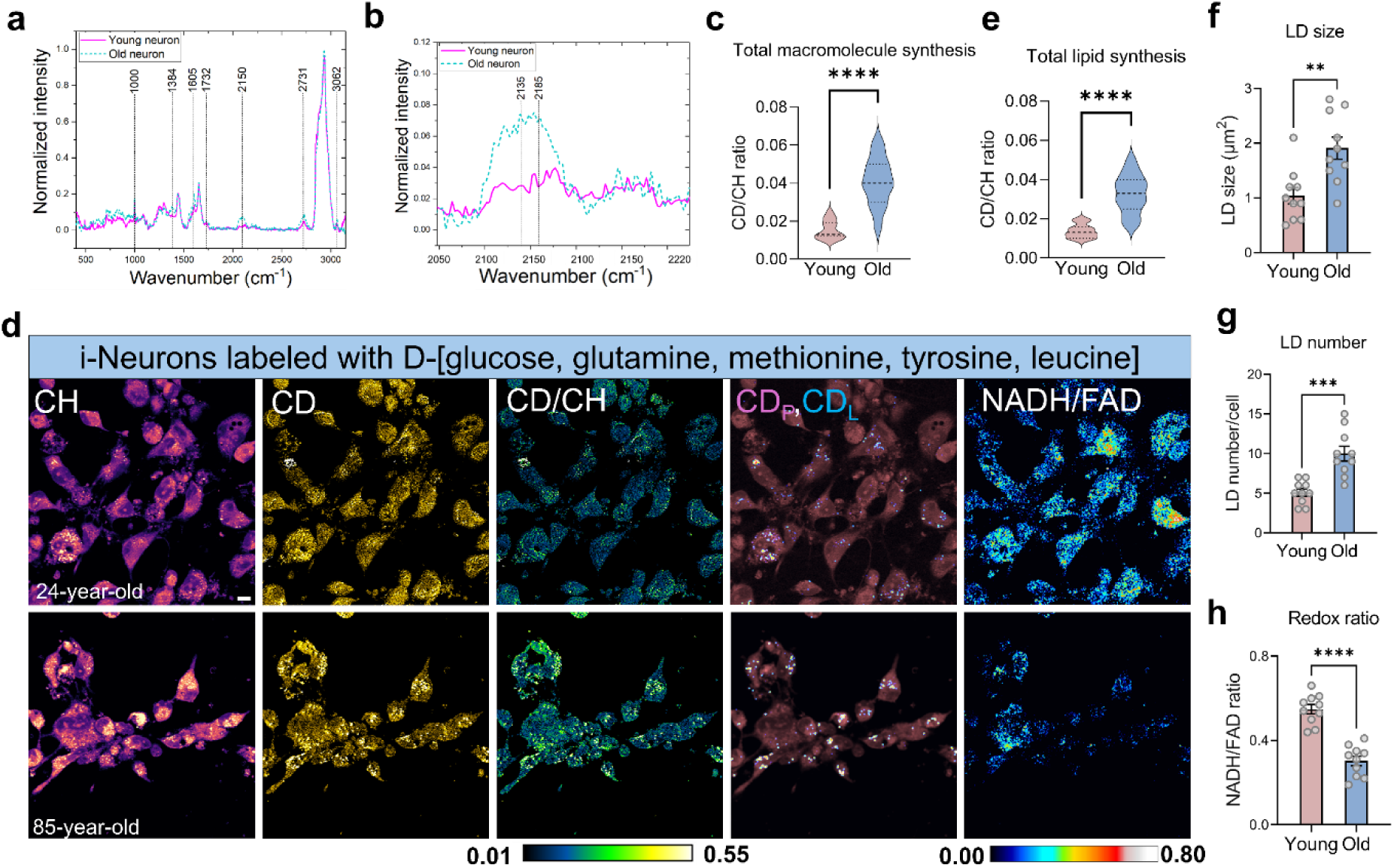
SuMMIT-SRS imaging of metabolic disorders in aging i-neurons. **(a)** Raman spectra of D-metabolites labeled young and old neurons. **(b)** Zoomed-in C–D region from (D) to show the shape and intensity differences of C–D spectra. **(c)** Quantification of the macromolecule synthesis by the ratio of CD (AUC) to CH (AUC) based on the Raman spectra in (D). n=15 spectra per group. **(d)** SRS imaging of C–H and C–D signal at 2932 and 2135 cm^-1^. The ratiometric images indicating global macromolecule turnover were generated by the intensity ratio CD/CH. The CD-protein (CD_P_) and lipids (CD_L_) were imaged and unmixed by SuMMIT-SRS. The NADH and FAD signal were captured by two photon excited fluorescent imaging, and the redox ratio was generated and shown as NADH/FAD. **(e)** Quantification of lipid synthesis in young and old neurons using the CD/CH ratio based on the SRS imaging shown in (d). **(f)** Quantification of lipid droplet size in young and old neurons based on the SRS imaging shown in (d). **(g)** Quantification of lipid droplet number in young and old neurons based on the SRS imaging shown in (d). **(h)** Quantification of redox ratio (NADH/FAD) in young and old neurons based on the SRS imaging shown in (d). Values in (c, e-h) are mean ± SEM. ns, not significant; **, p < 0.01; ***, p < 0.001; ****, p < 0.0001 by Student’s *t*-test.

## Notes

### Competing Interest Statement

The authors have declared no competing interest.

## References

1 Wei, Q., Qian, Y., Yu, J. & Wong, C. C. Metabolic rewiring in the promotion of cancer metastasis: mechanisms and therapeutic implications. Oncogene 39, 6139–6156, doi:10.1038/s41388-020-01432-7 (2020).

2 Chapman, N. M. & Chi, H. Metabolic rewiring and communication in cancer immunity. Cell Chem Biol 31, 862–883, doi:10.1016/j.chembiol.2024.02.001 (2024).

3 Motori, E. et al. Neuronal metabolic rewiring promotes resilience to neurodegeneration caused by mitochondrial dysfunction. Sci Adv 6, eaba8271, doi:10.1126/sciadv.aba8271 (2020).

4 Ma, S., Ming, Y., Wu, J. & Cui, G. Cellular metabolism regulates the differentiation and function of T-cell subsets. Cell Mol Immunol 21, 419–435, doi:10.1038/s41423-024-01148-8 (2024).

5 Pavlova, N. N., Zhu, J. & Thompson, C. B. The hallmarks of cancer metabolism: Still emerging. Cell Metab 34, 355–377, doi:10.1016/j.cmet.2022.01.007 (2022).

6 Drapela, S., Ilter, D. & Gomes, A. P. Metabolic reprogramming: a bridge between aging and tumorigenesis. Mol Oncol 16, 3295–3318, doi:10.1002/1878-0261.13261 (2022).

7 Plata-Gomez, A. B. & Ho, P. C. Age- and diet-instructed metabolic rewiring of the tumor-immune microenvironment. J Exp Med 222, doi:10.1084/jem.20241102 (2025).

8 Liu, L. et al. Metabolic reprogramming in T cell senescence: a novel strategy for cancer immunotherapy. Cell Death Discov 11, 161, doi:10.1038/s41420-025-02468-y (2025).

9 Guo, J. et al. Aging and aging-related diseases: from molecular mechanisms to interventions and treatments. Signal Transduct Target Ther 7, 391, doi:10.1038/s41392-022-01251-0 (2022).

10 Liu, H. et al. Energy metabolism in health and diseases. Signal Transduct Target Ther 10, 69, doi:10.1038/s41392-025-02141-x (2025).

11 Chen, X. et al. Metabolic Reprogramming Induced by Aging Modifies the Tumor Microenvironment. Cells 13, doi:10.3390/cells13201721 (2024).

12 Amorim, J. A. et al. Mitochondrial and metabolic dysfunction in ageing and age-related diseases. Nat Rev Endocrinol 18, 243–258, doi:10.1038/s41574-021-00626-7 (2022).

13 Sharifi, S. et al. Reducing the metabolic burden of rRNA synthesis promotes healthy longevity in Caenorhabditis elegans. Nat Commun 15, 1702, doi:10.1038/s41467-024-46037-w (2024).

14 Khalaf, F., Barayan, D., Saldanha, S. & Jeschke, M. G. Metabolaging: a new geroscience perspective linking aging pathologies and metabolic dysfunction. Metabolism 166, 156158, doi:10.1016/j.metabol.2025.156158 (2025).

15 Trefely, S. et al. Subcellular metabolic pathway kinetics are revealed by correcting for artifactual post harvest metabolism. Mol Metab 30, 61–71, doi:10.1016/j.molmet.2019.09.004 (2019).

16 Lee, W. D., Mukha, D., Aizenshtein, E. & Shlomi, T. Spatial-fluxomics provides a subcellular-compartmentalized view of reductive glutamine metabolism in cancer cells. Nat Commun 10, 1351, doi:10.1038/s41467-019-09352-1 (2019).

17 Chen, B., Lyssiotis, C. A. & Shah, Y. M. Mitochondria-organelle crosstalk in establishing compartmentalized metabolic homeostasis. Mol Cell 85, 1487–1508, doi:10.1016/j.molcel.2025.03.003 (2025).

18 Wei, L., Yu, Y., Shen, Y., Wang, M. C. & Min, W. Vibrational imaging of newly synthesized proteins in live cells by stimulated Raman scattering microscopy. Proc Natl Acad Sci U S A 110, 11226–11231, doi:10.1073/pnas.1303768110 (2013).

19 Wei, L. et al. Imaging complex protein metabolism in live organisms by stimulated Raman scattering microscopy with isotope labeling. ACS Chem Biol 10, 901–908, doi:10.1021/cb500787b (2015).

20 Shi, L. et al. Optical imaging of metabolic dynamics in animals. Nat Commun 9, 2995, doi:10.1038/s41467-018-05401-3 (2018).

21 Miao, K. & Wei, L. Live-Cell Imaging and Quantification of PolyQ Aggregates by Stimulated Raman Scattering of Selective Deuterium Labeling. ACS Cent Sci 6, 478–486, doi:10.1021/acscentsci.9b01196 (2020).

22 Li, Y. et al. Microglial lipid droplet accumulation in tauopathy brain is regulated by neuronal AMPK. Cell Metab, doi:10.1016/j.cmet.2024.03.014 (2024).

23 Li, Y. et al. Bioorthogonal Stimulated Raman Scattering Imaging Uncovers Lipid Metabolic Dynamics in Drosophila Brain During Aging. GEN Biotechnol 2, 247–261, doi:10.1089/genbio.2023.0017 (2023).

24 Zhang, L. et al. Spectral tracing of deuterium for imaging glucose metabolism. Nat Biomed Eng 3, 402–413, doi:10.1038/s41551-019-0393-4 (2019).

25 Li, X. et al. Quantitative Imaging of Lipid Synthesis and Lipolysis Dynamics in Caenorhabditis elegans by Stimulated Raman Scattering Microscopy. Anal Chem 91, 2279–2287, doi:10.1021/acs.analchem.8b04875 (2019).

26 Lu, F. K. et al. Label-free DNA imaging in vivo with stimulated Raman scattering microscopy. Proc Natl Acad Sci U S A 112, 11624–11629, doi:10.1073/pnas.1515121112 (2015).

27 Venkatesan, M. et al. Spatial subcellular organelle networks in single cells. Sci Rep 13, 5374, doi:10.1038/s41598-023-32474-y (2023).

28 Park, J. et al. Spatial omics technologies at multimodal and single cell/subcellular level. Genome Biol 23, 256, doi:10.1186/s13059-022-02824-6 (2022).

29 Becker, L. et al. Raman Imaging and Fluorescence Lifetime Imaging Microscopy for Diagnosis of Cancer State and Metabolic Monitoring. Cancers (Basel) 13, doi:10.3390/cancers13225682 (2021).

30 Zhu, A., Lee, D. & Shim, H. Metabolic positron emission tomography imaging in cancer detection and therapy response. Semin Oncol 38, 55–69, doi:10.1053/j.seminoncol.2010.11.012 (2011).

31 Emwas, A. H. et al. NMR Spectroscopy for Metabolomics Research. Metabolites 9, doi:10.3390/metabo9070123 (2019).

32 Chadwick, G. L., Jimenez Otero, F., Gralnick, J. A., Bond, D. R. & Orphan, V. J. NanoSIMS imaging reveals metabolic stratification within current-producing biofilms. Proc Natl Acad Sci U S A 116, 20716–20724, doi:10.1073/pnas.1912498116 (2019).

33 Zhang, D. S. et al. Multi-isotope imaging mass spectrometry reveals slow protein turnover in hair-cell stereocilia. Nature 481, 520–524, doi:10.1038/nature10745 (2012).

34 Steinhauser, M. L. et al. Multi-isotope imaging mass spectrometry quantifies stem cell division and metabolism. Nature 481, 516–519, doi:10.1038/nature10734 (2012).

35 Choe, M. & Titov, D. V. Genetically encoded tools for measuring and manipulating metabolism. Nat Chem Biol 18, 451–460, doi:10.1038/s41589-022-01012-8 (2022).

36 Qian, Y., Celiker, O. T., Wang, Z., Guner-Ataman, B. & Boyden, E. S. Temporally multiplexed imaging of dynamic signaling networks in living cells. Cell 186, 5656–5672 e5621, doi:10.1016/j.cell.2023.11.010 (2023).

37 Hu, F., Shi, L. & Min, W. Biological imaging of chemical bonds by stimulated Raman scattering microscopy. Nat Methods 16, 830–842, doi:10.1038/s41592-019-0538-0 (2019).

38 Saar, B. G. et al. Video-rate molecular imaging in vivo with stimulated Raman scattering. Science 330, 1368–1370, doi:10.1126/science.1197236 (2010).

39 Cheng, J. X. & Xie, X. S. Vibrational spectroscopic imaging of living systems: An emerging platform for biology and medicine. Science 350, aaa8870, doi:10.1126/science.aaa8870 (2015).

40 Freudiger, C. W. et al. Label-free biomedical imaging with high sensitivity by stimulated Raman scattering microscopy. Science 322, 1857–1861, doi:10.1126/science.1165758 (2008).

41 Jang, H., Wu, S., Li, Y., Li, Z. & Shi, L. Metabolic nanoscopy enhanced by experimental and computational approaches. Npj Imaging 2, 55, doi:10.1038/s44303-024-00062-y (2024).

42 Zhang, W. et al. Multi-molecular hyperspectral PRM-SRS microscopy. Nat Commun 15, 1599, doi:10.1038/s41467-024-45576-6 (2024).

43 Zhang, J. et al. 13C isotope-assisted methods for quantifying glutamine metabolism in cancer cells. Methods Enzymol 542, 369–389, doi:10.1016/B978-0-12-416618-9.00019-4 (2014).

44 Oh, S. et al. Protein and lipid mass concentration measurement in tissues by stimulated Raman scattering microscopy. Proc Natl Acad Sci U S A 119, e2117938119, doi:10.1073/pnas.2117938119 (2022).

45 Velez, D. O. et al. 3D collagen architecture induces a conserved migratory and transcriptional response linked to vasculogenic mimicry. Nat Commun 8, 1651, doi:10.1038/s41467-017-01556-7 (2017).

46 Chen, K. et al. Phenotypically supervised single-cell sequencing parses within-cell-type heterogeneity. iScience 24, 101991, doi:10.1016/j.isci.2020.101991 (2021).

47 Leineweber, W. D. et al. Divergent iron regulatory states contribute to heterogeneity in breast cancer aggressiveness. iScience 27, 110661, doi:10.1016/j.isci.2024.110661 (2024).

48 Ducker, G. S. & Rabinowitz, J. D. One-Carbon Metabolism in Health and Disease. Cell Metab 25, 27–42, doi:10.1016/j.cmet.2016.08.009 (2017).

49 Piper, M. D. et al. A holidic medium for Drosophila melanogaster. Nat Methods 11, 100–105, doi:10.1038/nmeth.2731 (2014).

50 Musselman, L. P. & Kuhnlein, R. P. Drosophila as a model to study obesity and metabolic disease. J Exp Biol 221, doi:10.1242/jeb.163881 (2018).

51 Beller, M. et al. PERILIPIN-dependent control of lipid droplet structure and fat storage in Drosophila. Cell Metab 12, 521–532, doi:10.1016/j.cmet.2010.10.001 (2010).

52 Liao, Y. et al. Amino acid is a major carbon source for hepatic lipogenesis. Cell Metab 36, 2437–2448 e2438, doi:10.1016/j.cmet.2024.10.001 (2024).

53 Ben-Moshe, S. & Itzkovitz, S. Spatial heterogeneity in the mammalian liver. Nat Rev Gastroenterol Hepatol 16, 395–410, doi:10.1038/s41575-019-0134-x (2019).

54 Franklin, T. B., Saab, B. J. & Mansuy, I. M. Neural mechanisms of stress resilience and vulnerability. Neuron 75, 747–761, doi:10.1016/j.neuron.2012.08.016 (2012).

55 Castelli, V. et al. Neuronal Cells Rearrangement During Aging and Neurodegenerative Disease: Metabolism, Oxidative Stress and Organelles Dynamic. Front Mol Neurosci 12, 132, doi:10.3389/fnmol.2019.00132 (2019).

56 Mertens, J., Reid, D., Lau, S., Kim, Y. & Gage, F. H. Aging in a Dish: iPSC-Derived and Directly Induced Neurons for Studying Brain Aging and Age-Related Neurodegenerative Diseases. Annu Rev Genet 52, 271–293, doi:10.1146/annurev-genet-120417-031534 (2018).

57 Rhine, K. et al. Neuronal aging causes mislocalization of splicing proteins and unchecked cellular stress. Nat Neurosci 28, 1174–1184, doi:10.1038/s41593-025-01952-z (2025).

58 Mertens, J. et al. Directly Reprogrammed Human Neurons Retain Aging-Associated Transcriptomic Signatures and Reveal Age-Related Nucleocytoplasmic Defects. Cell Stem Cell 17, 705–718, doi:10.1016/j.stem.2015.09.001 (2015).

59 Jang, H. et al. Super-resolution SRS microscopy with A-PoD. Nat Methods 20, 448–458, doi:10.1038/s41592-023-01779-1 (2023).

60 Homem, C. C. F. et al. Ecdysone and mediator change energy metabolism to terminate proliferation in Drosophila neural stem cells. Cell 158, 874–888, doi:10.1016/j.cell.2014.06.024 (2014).

61 Chell, J. M. & Brand, A. H. Nutrition-responsive glia control exit of neural stem cells from quiescence. Cell 143, 1161–1173, doi:10.1016/j.cell.2010.12.007 (2010).

62 Zhang, X. et al. Integrated multi-omics analysis of metabolome and transcriptome profiles during bovine adipocyte differentiation reveals functional divergence of FADS2 isoforms in lipid metabolism regulation. BMC Genomics 26, 457, doi:10.1186/s12864-025-11650-6 (2025).

63 Crotta Asis, A., Asaro, A. & D’Angelo, G. Single cell lipid biology. Trends Cell Biol, doi:10.1016/j.tcb.2024.12.002 (2025).

